# Habitat-specific environmental characteristics are associated with the movement of male and female loggerhead sea turtles

**DOI:** 10.64898/2026.05.04.722703

**Authors:** Perla Roman-Torres, Gail Schofield, Victor A. Stiebens, Christian Roder, Thomas Reischig, Herculano A. Dinis, Sandra Correia, Albert Taxonera, Graeme C. Hays, Christophe Eizaguirre

**Affiliations:** Queen Mary University of London, School of Biological and Chemical Sciences, Mile End Road E14NS, London, United Kingdom; Fundação Tartaruga, Riba D’Olte, Sal Rei, Cabo Verde; Associação Projeto Vitó, São Filipe, Fogo, Cabo Verde; Instituto do Mar, I.P. Cova de Inglesa, CP132, Mindelo, São Vicente, Cabo Verde; Associação Projeto Biodiversidade, Santa Maria, Sal, Cabo Verde; Deakin Marine Research and Innovation Centre, School of Life and Environmental Sciences, Deakin University, Geelong, VIC, Australia

**Keywords:** Movement Ecology, loggerhead sea turtle, habitat match, remigration interval, breeding periodicity, phenology

## Abstract

Linking animal movements to environmental drivers is essential for understanding ecological processes and anticipating species responses to climate change. We investigated habitat-specific movements in a globally significant aggregation of loggerhead turtles (*Caretta caretta*) nesting in Cabo Verde. Satellite tags on 15 adults (12 females, 3 males) provided multi-year tracks spanning breeding, migration, and foraging habitats. Movements and phenology differed by habitat. During the breeding season, females used either coastal areas, remaining within ∼20 m depth, or undertook long looping forays up to 360 km. Males showed two strategies: two remained resident in Cabo Verde waters, including Fra, the largest male tracked (Curved carapace length of 105 cm compared with a male mean of 90.7 ± 10.3 cm), while the third migrated annually to distant foraging grounds and returned ahead of the subsequent breeding season. In foraging habitats, turtles adopted neritic or oceanic strategies: neritic turtles remained localised in warm, productive waters, whereas oceanic turtles ranged widely in deeper, less productive areas. Time- and space-shift analyses showed that oceanic foragers used intermediate sea surface temperature and chlorophyll-a conditions relative to nearby or temporally shifted alternatives, consistent with movement within a thermal–trophic trade-off. Together, these results show how sex, body size, and energy balance drive habitat-specific movement dynamics in a changing ocean.

## Introduction

Understanding the factors that determine animal behaviour is a central goal of ecological, evolutionary, and conservation research (Block et al. 2011, Hussey et al. 2015, Hays et al. 2016, Wilson et al. 2019). Behaviour emerges from interactions between environmental drivers and adaptive processes, leading to habitat-specific patterns (Brown 1999, Sims et al. 2008, Habary et al. 2016, Holtmann et al. 2017, Camacho et al. 2020). Movements link directly to food acquisition, reproduction through conspecific interactions, and survival through interspecific interactions, thereby influencing species distributions and community structure (Nagelkerken and Munday 2015, Schofield et al. 2019, Rodríguez-Malagón et al. 2020). Yet, the environmental determinants of habitat-specific movements remain poorly understood, particularly in marine species with complex life cycles (Hays et al. 2016).

Studying the movement of marine vertebrates is challenging due to the three-dimensional nature of the oceans, and the large-scale distributions than can span entire ocean basins (Wallace et al. 2010, Block et al. 2011, Brodie et al. 2018). The development of technologies to remotely track marine megafauna in the marine environment has revealed that species’ distributions correlate with various environmental characteristics, including sea temperature, chlorophyll-a, oxygen, and even magnetic field (Sims et al. 2006, Kobayashi et al. 2008, Humphries et al. 2010, Hussey et al. 2015). Tracking has also shown that marine animals use different environments at different life stages (Shillinger et al. 2012, Riekkola et al. 2018, Sequeira et al. 2018). For example, blue sharks (*Prionace glauca*) in the Northeast Pacific exhibit sex- and age-dependent foraging preferences linked to thermal tolerance, with seasonal overlap likely coinciding with mating (Maxwell et al. 2019).

Sea turtles also show sex- and age-dependent differences in habitat use (Schofield et al. 2009, González Carman et al. 2016, Scott et al. 2017). Females can be readily studied when they emerge to nest, while adults and juveniles are captured in coastal waters (Hatase et al. 2002, Schofield et al. 2007, Godley et al. 2008, Marcovaldi et al. 2010, Hawkes et al. 2011, Pikesley et al. 2014). During breeding, females typically remain coastal, resting in near-shore areas within 20 m depth, directly in front of nesting beaches or within a few kilometres (Mortimer and Portier 1989, Schofield et al. 2009, Hart et al. 2010). These areas often provide sea surface temperatures suitable for egg maturation (Schofield et al. 2009, Weber et al. 2011). At cooler sites, females select wind-sheltered areas with warmer water to accelerate egg development (Schofield et al. 2010c, Fossette et al. 2012). While many females reduce movement during the nesting period, others undertake long forays between nesting events, sometimes hundreds of kilometres offshore. These strategies may balance access to thermal refugia and food resources that support egg maturation (Rees et al. 2010, Esteban et al. 2015).

While female movement behaviours have been relatively well characterised (Hatase et al. 2002, Schofield et al. 2007, Godley et al. 2008, Marcovaldi et al. 2010, Hawkes et al. 2011, Pikesley et al. 2014), male movements remain poorly documented because males rarely come ashore. Males appear to return more frequently than females to the mating grounds, likely to secure mating opportunities and balance operational sex ratios (Hays et al. 2014, Schofield et al. 2017). In some regions, some males can even remain resident in mating areas (Schofield et al. 2009, Arendt et al. 2011).

At the end of breeding, all females and some males migrate to foraging grounds, with males typically departing earlier (Van Dam et al. 2008, Schofield et al. 2009, Arendt et al. 2011). Migration tends to be directional and independent of currents (Plotkin 2003, Hays et al. 2014). Turtles remain at the foraging grounds until they build sufficient reserves to fuel the next reproductive migration (Plotkin 2003). Adults forage either oceanically or in coastal habitats (Hawkes et al. 2006, Hatase et al. 2007), with coastal foragers following diel activity cycles (Dujon et al. 2018) and oceanic individuals combining active swimming with drifting in response to thermal and productivity gradients (Willis-Norton et al. 2015, Polovina et al. 2017). Although displacement has been linked to temperature, no clear explanation exists for how turtles select different movement strategies across habitats.

Identifying these environmental drivers requires tracking individuals across breeding, migration, and foraging phases. The Cabo Verde archipelago offers an ideal setting, as its ten islands host distinct nesting groups exposed to different coastal environments (Stiebens et al. 2013, Baltazar-Soares et al. 2020). Earlier tracking showed that turtles from Cabo Verde forage either oceanically or in neritic waters off West Africa (Hawkes et al. 2006, Pikesley et al. 2014). Subsequent work extended this picture by showing that adult males from Boa Vista also use diverse movement strategies, including migration to oceanic and neritic West African waters and possible residency near the breeding area (Varo-Cruz et al. 2013). This work highlights how male movements remain under-sampled, while reinforcing the need to compare environmental drivers of movement among habitats, sexes and reproductive phases. Stable isotope studies further revealed that oceanic foragers split into two groups, one in upwelling areas and one in open ocean away from the continental shelf (Cameron et al. 2019). Oceanic turtles exploit vast regions in deep waters (>500 m), whereas neritic turtles travel to the continental shelf of Sierra Leone, where they remain in smaller areas within the 100 m isobath (Hawkes et al. 2006). Yet, we still need formal comparison of the environmental determinants of movement across habitats or between sexes in the face of climate-driven changes that may reshape individuals’ distributions. Here, we examined habitat-specific environmental parameters influencing movement in breeding, migratory, and foraging habitats using satellite telemetry and in-silico time- and space-shift experiments on 15 adult turtles.

## Methods

### Study area

The Republic of Cabo Verde is an archipelago composed of 10 volcanic islands and supports one of the largest aggregations of loggerhead sea turtles in the world (Taxonera 2022). This nesting aggregation is genetically distinct from all other nesting aggregations of the Atlantic (Reis et al. 2009, Monzón-Argüello et al. 2010, Baltazar-Soares et al. 2020), and is composed of independent genetic breeding groups of conservation value (Stiebens et al. 2013, Baltazar-Soares et al. 2020).

### Tag deployment

Between July and September of 2011-2013, 15 adult loggerhead sea turtles (12 females and 3 males) were equipped with satellite-relayed devices, including six Satellite Relay Data Loggers (SDRL) Fastloc GPS Argos devices manufactured by SMRU (Sea Mammal Research Unit, St. Andrews, UK; n_Boa Vista_ = 3, n_Sal_ = 3). In 2012, 2 Splash 203 from Wildlife computers (Redmond, WA, USA) and one Fastloc F4G 291 from Sirtrack (Havelock North, New Zealand) were deployed on females nesting on the beaches of São Vincente. In 2012, 1 Splash 203 and two Fastloc F4G 291 were deployed on turtles nesting on Fogo Island. Lastly, in 2013, three males from Boa Vista were equipped with a SPLASH10-295C transmitter from Wildlife computers (Table S1). Males were captured in water by fishermen, and females were stopped after they had completed nesting. Devices were secured with an epoxy glue (Pure-2K, Power fasteners, New Rochelle, NY, U.S.A., Schofield et al. 2010a). At the time of deployment, the curved carapace length (CCL) and the curved carapace width (CCW) were measured (±5 mm). All turtles were marked with passive integrated transponder tags to facilitate individual identification on their return to nest should they lose the device (Stiebens et al. 2013).

#### Data filtering

We used the Satellite Tracking and Analysis Tool (STAT) from www.seaturtle.org to save and manage data (Coyne and Godley 2005). We used “sdafilter” from the “Argosfilter” R package, which removed all locations of poor-quality class (class B and 0), those with less than 3 satellites, and turtles showing unrealistic swimming speed (>2 m/s) using McConnell et al.’s (1992) algorithm. This filter also removed all locations with impossible turning angles at the highest speed detected. This filtering resulted in a high-quality dataset composed of 14,213 locations (derived from Argos and Fastloc) for the 15 turtles (Table S1, S2).

### Defining movement patterns

Turtle movement phases were classified into breeding, migration, and foraging habitats based on location and behaviour. We defined the breeding habitats as the area used by the turtles in the Cabo Verde waters. The migratory route links Cabo Verde to the foraging habitat, offshore of Western Africa and on the coast of Sierra Leone (Hawkes et al. 2006, Pikesley et al. 2014). The foraging habitat refers to the place where turtles search for food either oceanically or neritically. After identifying the points of switch between phases (e.g. from breeding to migration to foraging), the average daily displacement (*D*_*i*_) was calculated per month as the distance travelled between successive positions divided by the number of days in the interval following Eq.1. The monthly total displacement is the total displacement between each point recorded for the month 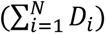.

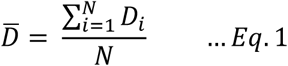

Directional and turning angles were derived following Equations 2–3, with angular metrics computed from consecutive positions.

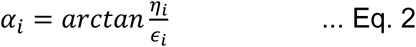

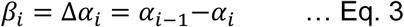

where α_i_ is the directional angle, *∈*_*i*_ is the longitude velocity, *η*_*i*_is the latitude velocity, *β*_*i*_ is the turning angle and Δ*⍺*_*i*_is the difference between the two angles. Because turning angles are sensitive to location error when successive positions are close in time or space, we interpreted these analyses as broad indicators of directional versus tortuous movement among habitats, rather than as fine-scale behavioural classifications.

### Environmental data acquisition

The concentration of chlorophyll-a (Chl-a, log transformed), sea surface temperature (SST) and bathymetry data were extracted using the “xtractomatic” R package (Mendelssohn 2008). Chl-a and SST data were extracted monthly (July 2011–June 2015, 0.05° resolution) from MODIS-Aqua via the R package *xtractomatic* (Mendelssohn, 2008), and bathymetry data were obtained from NOAA’s National Geophysical Data Center. These parameters were linked to each filtered turtle location to characterise the conditions experienced in different habitats.

### Time and space-shift experiments

To understand the environmental requirements of turtles specifically in oceanic foraging habitat, since this is the habitat in which they spend most of their time, we conducted two *in-silico* experiments, focusing Chl-a and SST.

In a time-shift experiment, we compared Chl-a and SST conditions experienced by turtles at their actual foraging locations with conditions one month earlier and one month later, simulating earlier or delayed arrival. A one-month interval ensured realistic but distinct environmental conditions and allows identifying potential mismatches between behaviours and environmental variables. Chl-a and SST were extracted similarly to the observed data.

A space-shift experiment tested whether turtles, following their exact track but in a different realistic location would be exposed to the same Chl-a and SST conditions as they did on a particular day. This approach enabled us to identify whether the environmental conditions rather than the location matters most. This virtual experiment consisted of creating a 3×3 identical polygon grid centred on the first observed location of the turtle in a day. We allowed the width of the polygons to vary daily, based on the distance travelled in a straight line by a given turtle between the first location of a day and the first location of the following day (Fig. S1A-C). This approach accounts for both inter-individual and intra-individual variations of daily habitat use and movement. We then extracted the mean concentration of Chl-a and SST within each polygon, and simulated a turtle choosing the highest and lowest levels. These pathways were then compared to the real pathways taken by turtles (Fig. S1D). This approach offers a statistical means of testing whether turtles i) maximize, ii) minimise, or iii) optimise their Chl-a and SST as the best environmental niche available.

For these two virtual experiments that focused on oceanic foraging habitat, we used the data of 10 individuals (nine females and one male) for which we received at least one daily location during the period of transmission. For both in-silico experiments, good quality Chl-a concentration and SST data were required. Therefore, for the time shift experiment, all months in which <70% Chl-a concentrations were recorded across the foraging area were removed (76.31% of data were retained). In the same way, for the spatial shift experiment, we removed all days where Chl-a values were not recorded at the locations of interest. If a turtle location remained within 10 km on a given day, the Chl-a image resolution was not sufficient, and these days were also removed. Ultimately, 898 days out of 1171 possible days were retained (76.68% of the possible data points).

### Statistical analyses

All statistics were conducted in R 3.6.3 software (R Core Team 2020).

#### Defining movement in the three habitats (breeding, foraging, migratory corridor)

We first characterised the movement of the turtles by testing whether turning angles differ from a uniform distribution using the Rayleigh test (rayleigh.test) in each habitat. We also performed an analysis of variance for circular data (aov.circular), to test differences in turning angle distribution in the three habitats using the R package “Circular”. We then compared the average daily displacement within each habitat using a linear mixed-effect model (LME, in packages “lme4”, and “lmertest”, “glht” in package “multicomp”), where turtle ID was set as a random factor to account for repeated measures.

To characterise the environmental attributes of each habitat, we tested whether the bathymetry, Chl-a concentration and SST experienced by the turtles differed using a LME with turtle ID set as a random effect, followed by a Tukey multicomponent test. Two turtles, Nhanha and Manga, were only included in the breeding habitat analysis, since Nhanha’s transmitter stopped emitting at 27 days post-tagging and Manga had low transmissions with few points per week till she stopped transmitting at 60 days post tagging (Table S1).

#### Within the breeding habitat

To distinguish movement differences during the nesting season, we tested the distribution of the turning angles of turtles breeding on different islands, using a Rayleigh test, and we tested turning angle distributions among island of origin and between sexes using circular ANOVA. We tested whether the average daily displacement, bathymetry, Chl-a concentration and SST varied with respect to island on which turtles nested and between sexes, using an LME with turtle ID set as a random effect. Furthermore, we tested whether bathymetry was correlated with Chl-a and SST using an LME with turtle ID set as a random effect.

#### Within the migratory corridors

To test possible differences between the neritic and oceanic turtles (9 females and one male) and between sexes (10 females and one male) during migration, we examined the average daily displacement and the experienced bathymetry, Chl-a and SST using an LME with turtle ID set as a random effect. We tested whether the average daily displacement was correlated with Chl-a and SST using an LME, with turtle ID set as a random effect.

#### Within the foraging habitat

To identify differences between neritic and oceanic foraging movement patterns, we tested respective turning angle distributions using the Rayleigh test. We also tested differences in turning angle distribution and average daily displacement between these two strategies using a circular ANOVA and an LME with turtle ID as a random effect, respectively. For the oceanic strategy, we tested differences between sexes (9 females and one male), for the turning angle, average daily displacement, experienced bathymetry, SST and Chl-a concentration using ANOVA for circular data and an LME, with turtle ID set as a random effect, respectively. For the neritic strategy, we tested differences between the resident males in Cabo Verde and the neritic female forager, for the turning angle, average daily displacement, experienced bathymetry, SST and Chl-a concentration using ANOVA for circular data and an LME, with turtle ID set as a random effect, respectively.

#### Spatial and time shift experiments

To test the truly experienced SST and Chl-a and the experienced at the two hypothetical scenarios, we used an LME with turtle ID set as a random effect. For the spatial shift experiment, we compared Chl-a concentration and SST in the polygon where a given turtle was located, using the maximum and minimum Chl-a and SST values in the polygons. For this, we used two LMEs with turtle ID set as a random factor, followed by a pairwise Tukey test.

## Results

### Defining movements across habitats

Loggerhead turtles exhibited movements between 8°N–22°N latitude and 26°W–16°W longitude (Fig. 1A). Fifteen adults (12 females, 3 males) were tracked for an average of 390 days (range: 27–1,109 days; Table S1). Mean daily displacement was 16.9 ± 6.8 km, with the longest total distance recorded for *Bemvinda* (17,073 km over 1,109 days). No sex differences were detected in daily displacement (LME F₁,₁₁ = 0.135, p = 0.727).

**Figure 1.**
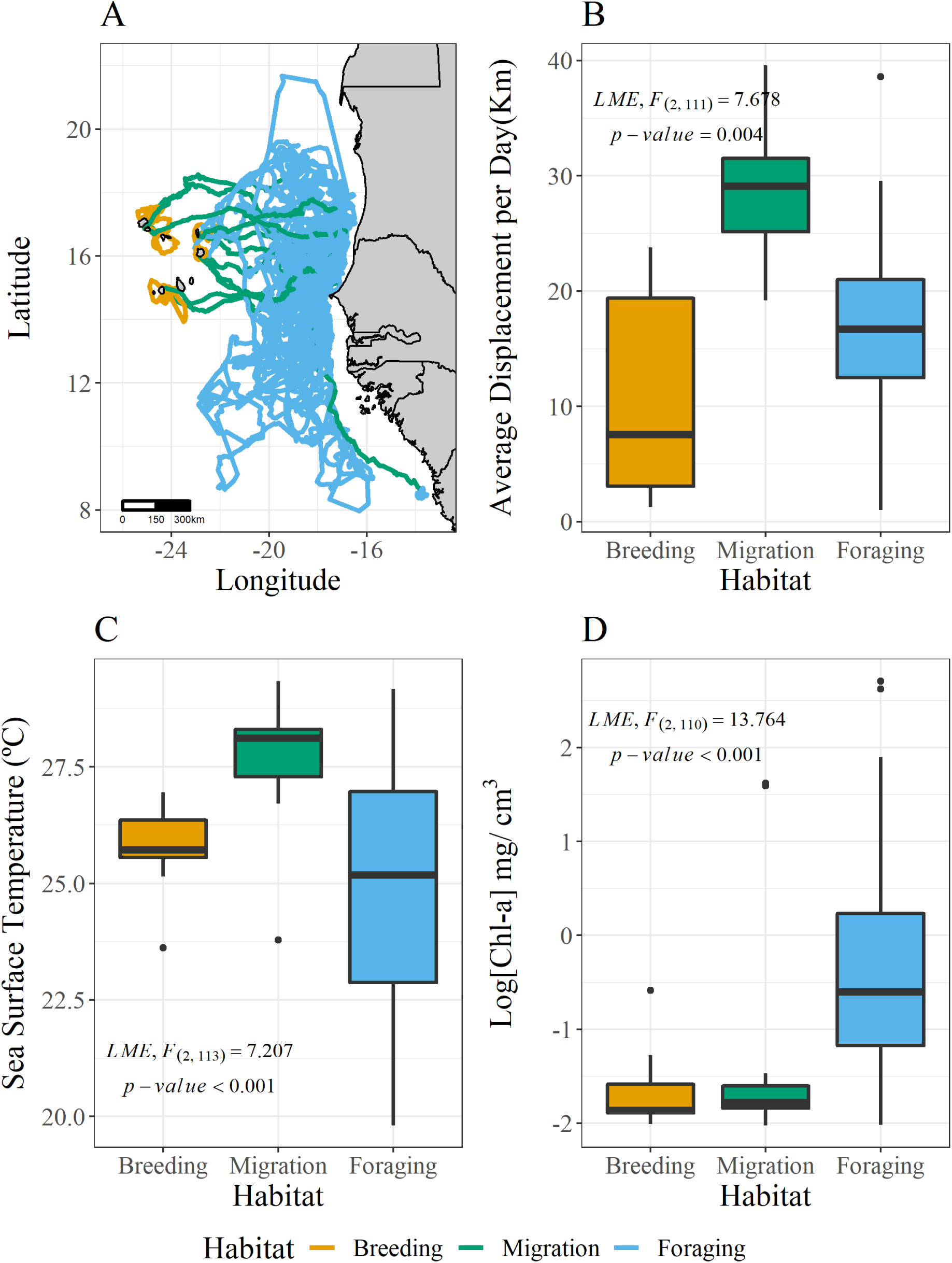
Turtle tracks in the breeding ground, migratory corridor and foraging habitat (A). turtles’ average daily displacement (B) concentration of chlorophyll-a (C) sea surface temperature (D) experienced in the three habitats.

Movement patterns differed significantly among habitats (Circular ANOVA F₂,₁₄₁₈₁ = 3.85, p = 0.021). Displacement was greatest during migration (28.3 ± 3.6 km day⁻¹), followed by foraging (17.3 ± 6.3 km day⁻¹) and breeding (10.3 ± 8.4 km day⁻¹; LME F₂,₁₁₁ = 7.68, p = 0.004; Fig. 1B, Table S3). Migration was also the most directional phase, characterised by smaller turning angles (Figs. S2–S5).

Environmental parameters varied among habitats (LME Bathymetry F₂ = 22.57; SST F₂ = 7.21; Chl-a F₂ = 13.76; all p < 0.001). Turtles experienced the shallowest waters during breeding (–819 ± 995 m), deeper waters during migration (–3,063 ± 394 m) and foraging (–2,916 ± 1,231 m). SST peaked during migration (28.1 ± 0.7 °C) compared with breeding (25.8 ± 0.9 °C) and foraging (24.8 ± 2.6 °C, Fig. 1C). Chl-a concentrations were highest in foraging areas (–0.39 ± 1.02 mg m⁻³), intermediate in migratory routes, and lowest during breeding (Fig. 1D). To better characterize habitat-specific movements, we then focused on each habitat type separately.

### Breeding habitat

#### Phenology of migration

Turtles remained in breeding areas from early July until departures that began in late July (males) and extended to early October (females, Fig. 2). Departure timing differed strongly between sexes: the male departed around Julian day #201 (July 20th), while females departed on average around Julian day #259 (∼mid-September) — an average difference of 58 days, consistent with males leaving significantly earlier in the season (Fig. 3A). This temporal separation likely reflects differences in reproductive investment and energy allocation between sexes.

**Figure 2.**
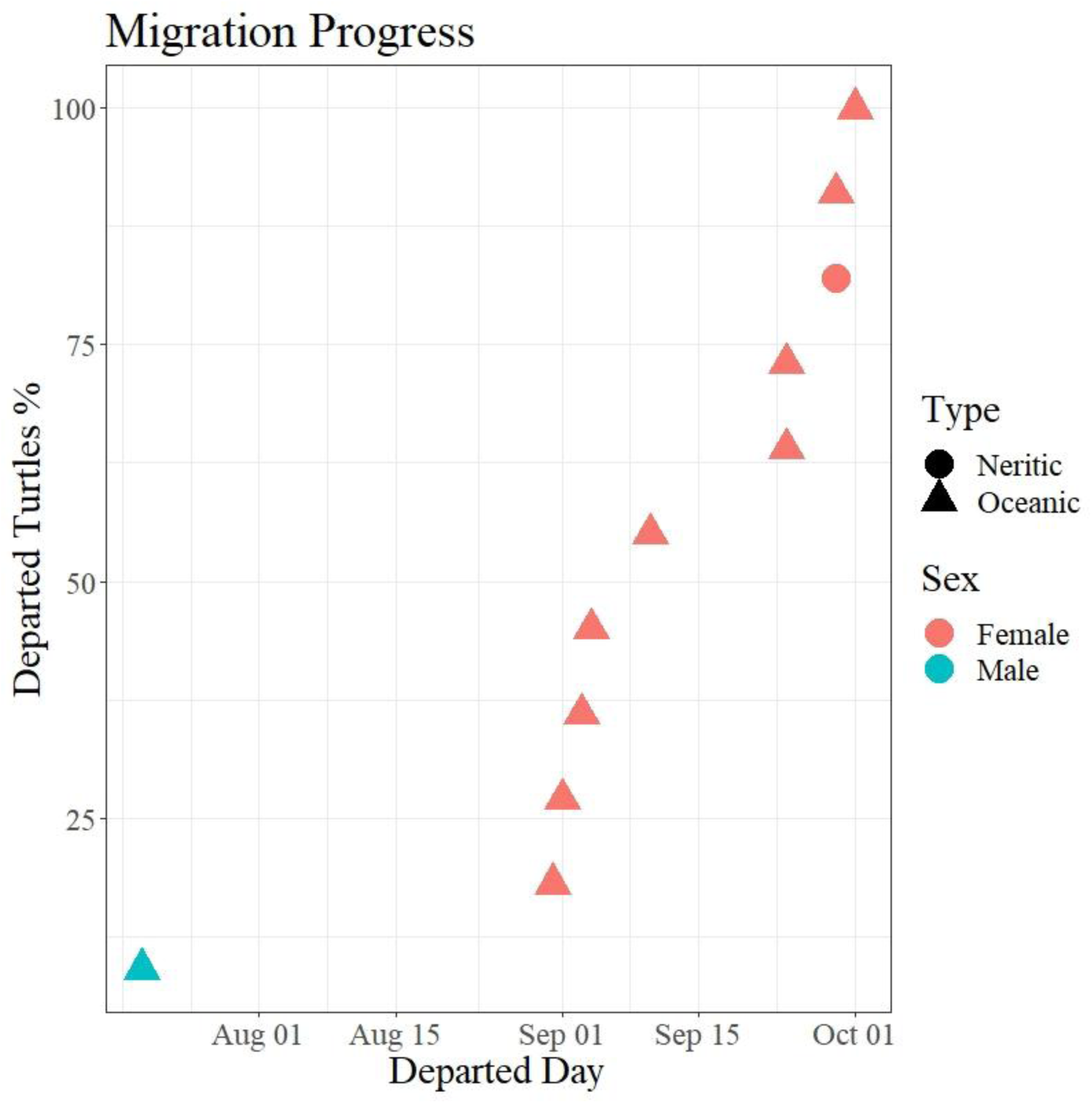
Proportion of turtles as a function of their departure time from the nesting ground. Both sexes are shown as well as their foraging strategy (neritic vs oceanic).

**Figure 3.**
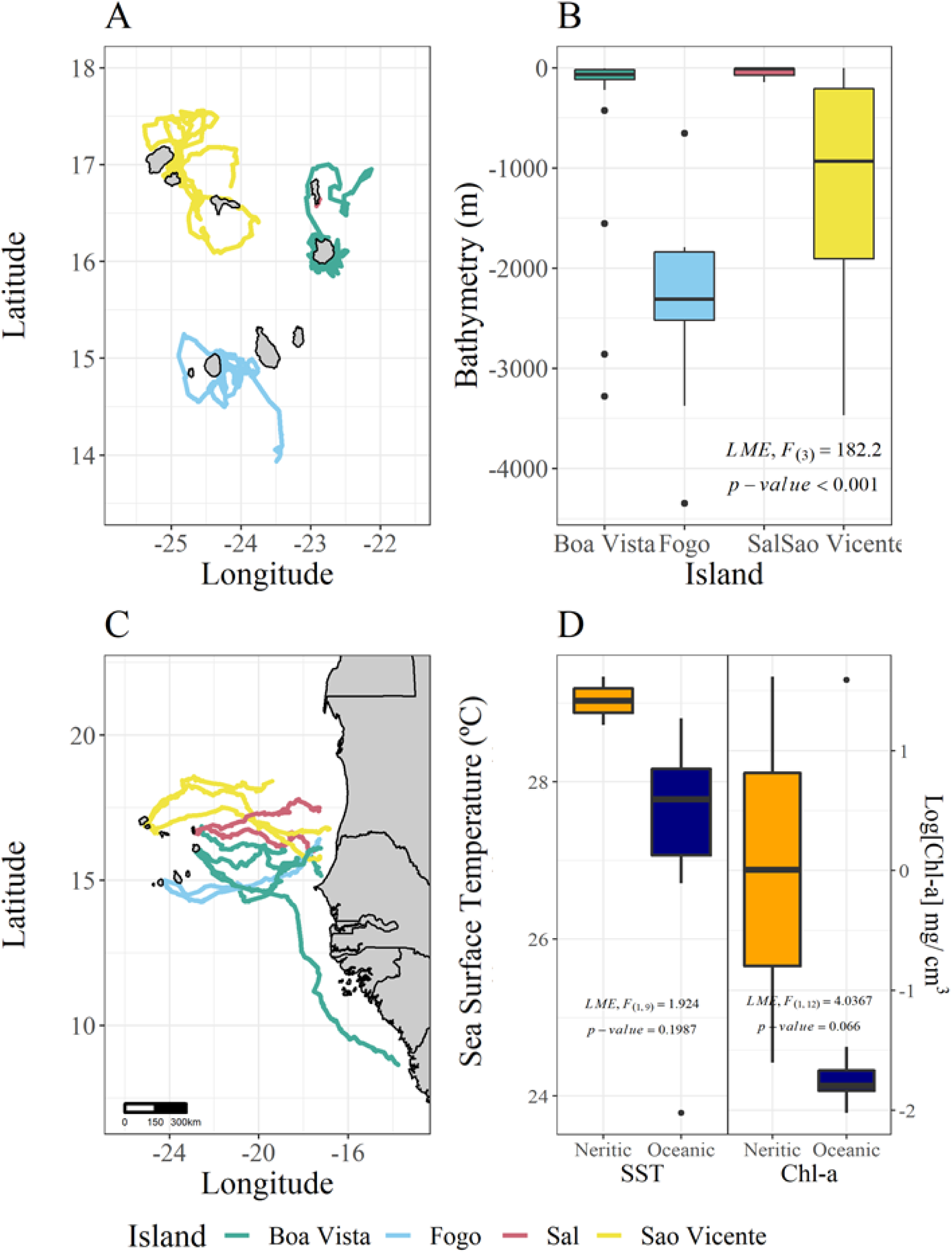
Movement within the breeding habitat. Individual tracks are coloured by breeding island (A). Bathymetry experienced by the turtles breeding on the four islands (B). In the migratory route, turtles follow a straight displacement towards Mauritania (C). Concentration of chlorophyll-a and sea surface temperature experienced by the neritic and oceanic turtles in their migratory route (D)

#### Movement patterns and habitat use

Females and males showed no difference in their average daily displacement and in the distribution of the turning angles (LME average daily displacement F_(1,11)_= 0.215, p= 0.651; Circular ANOVA, F_(1,1)_= 447.1, p<0.001). Males and females used broadly similar environmental conditions within the breeding habitat, with no detectable sex differences in bathymetry, SST or Chl-a (LME Bathymetry, F_(1,11)_= 0, p= 0.999; LME SST, F_(1,11)_= 1.792, p= 0.214; LME Chl-a, F_(1,11)_= 0.699, p= 0.427; Table S4). Among females, six individuals displayed long forays extending offshore, while six remained coastal (Fig. 3A). Oceanic turtles were from Fogo and São Vicente and showed a directional distribution in the turning angles (Rayleigh test, Test _Fogo_= 0.457, p<0.533; Test _Sal_= 0.347, p<0.001; Fig. S2), performing wide-ranging loops around the islands between nesting events, being exposed to deep bathymetry (Fig 3B). At the beginning of the foray, females swam against the currents for 7 to 9 days, before turning and making their way back following the current’s direction. In contrast, turtles nesting on Boa Vista and Sal presented a uniform distribution for turning angles (Rayleigh test, Test _Boa Vista_= 0.095, p= 0.533; Test _Sal_= 0.085, p<0.001; Fig. S3), displaced much less (F_(3,10)_=5.73, p =0.015) and remained in shallow coastal waters (Bathymetry, LME, F_(3,10)_=182.2, p <0.001, Table 4, Fig. 3B). As expected from those observations, the average daily displacement was negatively correlated with the bathymetry experienced by turtles (LME, Daily displacement = -0.007 Bathymetry + 7.541, Pearson Correlation test, t_(1,31)_= -6.722, p<0.001, R^2^= -0.888).

Out of the three males, two were residents (Fra and Ze) in the sense that they remained during the entire tag transmission period in Cabo Verde waters, in the south-west coasts of Boa Vista, overlapping partially with the females’ distribution (Fig. S6). During this time, resident males had similar movement patterns to females nesting at Sal and Boa Vista (Table S4). At the end of the breeding period, Fra, the largest males (curved carapace length 105 cm, mean_male_ = 90.7 ± 10.3 cm) remained in this area, while Ze explored the northeast of the island. The third male, Mingo, migrated away from Boa Vista, swam north around Sal island, and left Cabo Verde toward the foraging area at the end of July. This male had a similar swimming pattern to females from Fogo and São Vicente (Table S4). In the breeding areas, neither Chl-a concentrations (LME F_(3,10)_= 0.230, p= 0.867) nor SST showed variation across islands (LME, F_(3,10)_= 1.234, p= 0.348). It is therefore not surprising that the average daily displacement did not correlate with productivity (Chl-a, LME, F_(3,10)_= 0.230, p= 0.866) or temperature (SST, LME, F_(1,91)_=0.060, p=0.808, R^2^=0.247).

#### Migratory corridors

Migration began earlier in males (late July) than in females (September–October). The male and nine females moved eastward toward Mauritania and Senegal before initiating foraging in deep pelagic waters (bathymetry < –3,000 m), while one female reached Sierra Leone’s neritic shelf (bathymetry > –50 m). Migration was highly directional (Rayleigh F = 0.36, p < 0.001, Fig. 2C), with no sex or strategy differences (Circular ANOVA p > 0.8). Mean displacement was 29.7 ± 4.7 km day⁻¹ in females and 22.2 ± 4.2 km day⁻¹ in the male (LME F₁,₁₀ = 3.08, p = 0.130, Fig. S4).

Bathymetry and Chl-a concentrations experienced during migration differed slightly between migrations to oceanic and neritic areas, with higher productivity and lower bathymetry in the neritic zone (LME _Bathymetry_, F_(1,11)_=5.166, p= 0.044; Bathymetry _Neritic_= 1747 SE ± 2329, Bathymetry _Oceanic_= -3147 SE ± 399; LME _[Chl-a],_ F_(1,11)_=4.036, p=0.067, [Chl-a] _Neritic_ = 0.008 mg m^−3^ SE ± 2.27, [Chl-a]_Oceanic_ = -1.417 mg m^−3^ SE ± 1.138; Fig. 2D). There was no difference in terms of temperature (LME SST, F_(1,11)_= 1.924, p= 0.198, SST _Neritic_=29.03°C SE ± 0.44, SST _Oceanic_ =27.90°C SE ±0.59). There was no difference between sexes for either of the environmental variables investigated (LME _Bathymetry,_ F_(1,11)_= 0.910, p= 0.361 Bathymetry _Females_= 2823 SE ± 974, Bathymetry _Male_= 3506 SE ±175; LME _[Chl-a],_ F_(1,10)_= 0.332, p= 0.575, [Chl-a] _Females_ = -1.158 SE+-1.374; [Chl-a] _Male_ = -1.739 SE ± 0.139), except for temperature where females were exposed to higher temperature on average than males (LME SST, F_(1,10)_= 16.349, p= 0.001, SST _Females_ =28.107 SE ± 0.710, SST _Male_ =25.285 SE ± 2.066). During migration, the average daily displacement did not correlate with any environmental variable (LME_CHL-A_, F_(1,2)_=1.836, P=0.288; LME_SST_, F_(1,5)_=3.172, P=0.127).

#### Foraging habitat

Of ten females with sufficient data, nine adopted oceanic and one neritic feeding strategy (Fig. 4 A-B). The neritic forager was from Boa Vista. We also confirmed that males exhibited both neritic (2 resident males to Boa Vista) and oceanic (1 migratory male) foraging strategies (Fig. 4 C-D), however, in this case, neritic males did not migrate to Sierra Leone but remained in the shallow waters of Cabo Verde.

**Figure 4.**
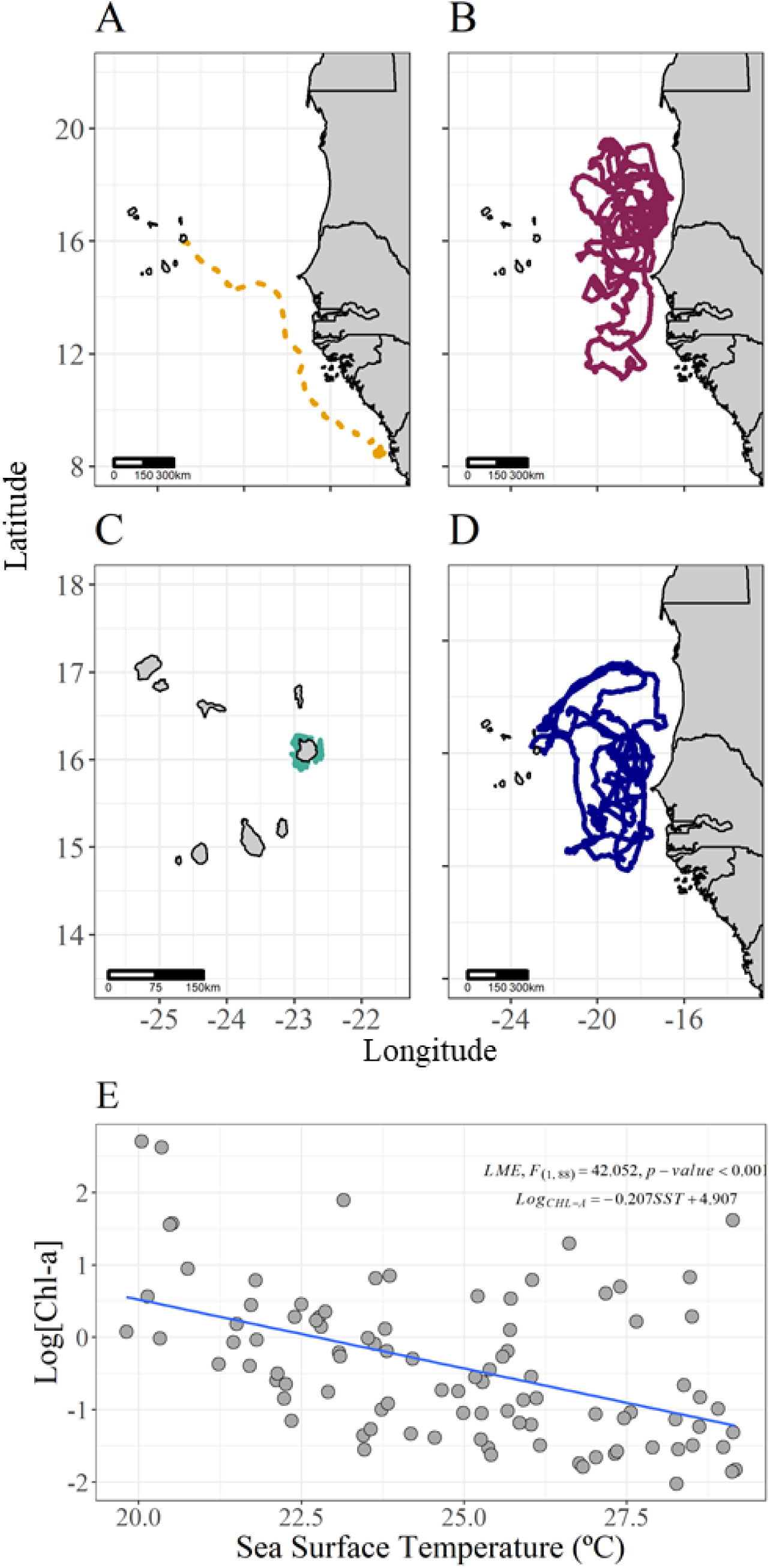
Within the foraging, neritic (A) and oceanic (B) female turtle tracks are shown. Neritic (C) and oceanic (D) male turtle tracks. Correlation between chlorophyll concentration and sea surface temperature in the foraging habitat (E).

Combining females and males, we found the average daily displacement of neritic turtles (one female and two resident males) was significantly shorter than that of oceanic turtles (Nine females and migratory male) (Displacement of oceanic turtles = 17.8 (SE± 6.1) km day⁻¹; Neritic turtles = 9.1 (SE ± 2.4) km day⁻¹. LME, F_(1,10)_= 8.569, p=0.018). Among oceanic foragers, *Olympia* and *Nusco* travelled furthest (25.9 ± 1.5 and 22.9 ± 4.0 km day⁻¹, respectively). Displacement did not correlate with SST or Chl-a (LME_CHL-A_, F_(1,92)_=0.094, P=0.760; LME_SST_, F_(1,90)_=0.870, p=0.354).

No sex differences were found for turning angles, displacement, bathymetry, SST, or Chl-a among oceanic foragers (all p > 0.2), Table S4), suggesting they use similar environmental niches. Similarly, neritic individuals showed no differences between sexes for most variables, except for productivity: females experienced higher Chl-a (0.93 ± 0.44 mg m⁻³) than males (–0.61 ± 0.73 mg m⁻³; LME F₁,₂ = 20.0, p = 0.0009, Table S3), likely because of their location in Western Africa.

The single neritic female remained in a highly productive 40 km² shelf area (Chl-a = 2.75 ± 0.12 mg m⁻³; SST = 27.1 ± 1.5 °C), moving ∼9 km day⁻¹ with a uniform turning angle distribution (*p*=0.28). These values contrasted significantly with oceanic foragers (displacement F₁,₁₁ = 8.57, p = 0.018; Circular ANOVA F₁,₁ = 98.0, p < 0.001, Fig. S5).

Oceanic turtles (nine females and one migratory male) over deep waters, beyond the continental shelf (Average bathymetry= -3110 m isobath SE ± 881m, LME _Bathymetry_, F_(1,10)_= 12.04, p=0.008) with an average daily displacement of 17.81 ± 4.35 km day^−1^. Turning angle showed a distribution trend toward an excess of small angles (Rayleigh test, t_(9749)_=0.367, p<0.01, Fig. S5 B-K). Primary production in the oceanic area was significantly lower than in the neritic area with a Chl-a concentration of 0.995 mg m^−3^ ± 0.18 (LME _[Chl-a]_, F_(1,10)_= 6.162, p = 0.032). The sea surface temperature was not different in the oceanic and neritic habitat (LME _SST_, F_(1,16)_= 2.224, p=0.115).

#### Habitat use

Overall, our space and time-shift experiments showed that turtle movement in oceanic foraging areas is constrained by a negative correlation between sea surface temperature and chlorophyll-a concentration across time and space (LME, F_(1,88)_= 42.052, p<0.001; Fig. 4E). In the time-shift experiment, we show that Chl-a would have been lower if turtles had arrived one month earlier and similar if they had arrived one month later (LME F₃ = 9.18, p < 0.001; Fig. 5A, C). Conversely, SST was warmer one month earlier and cooler one month later (F₃,₂₇₆ = 27.45, p < 0.001; Fig. 5B, D, Table S6). Thus, earlier foraging exposed turtles to warmer, less productive waters, while later foraging provided cooler, more productive conditions.

**Figure 5.**
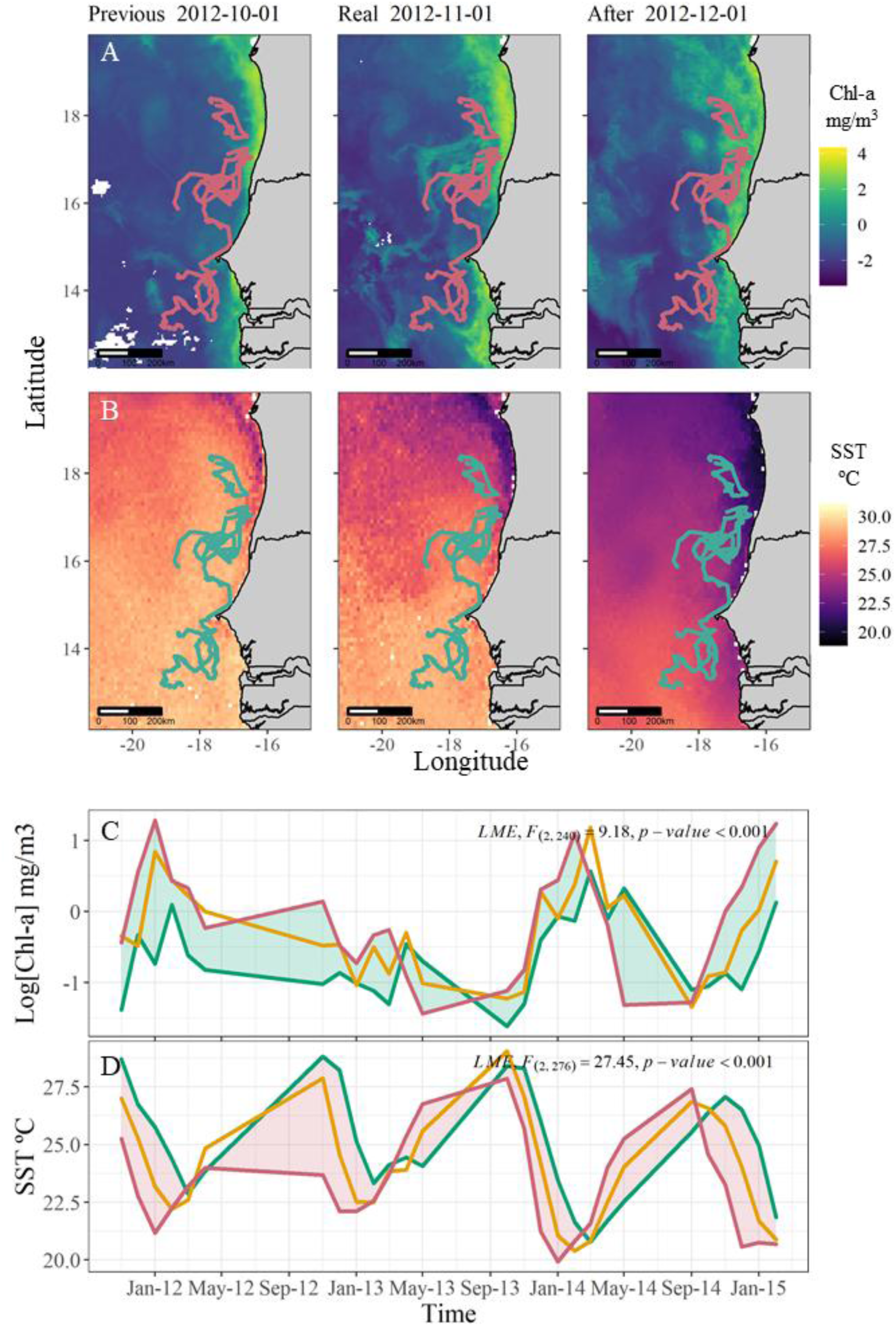
Time shift experiments. (A) specifically focuses on November 2012, when most turtles transmitted simultaneously as an example. The central map shows the observed concentrations of chlorophyll-a experienced by the turtles in November 2012. The left panel shows the concentrations that turtles would have experienced one month earlier. The right panel shows the concentrations turtles would have experienced if they had migrated one month later. (B) shows the results of the time shift experiment for sea surface temperature in November 2012, when most transmitters were active. (C) shows the results of the time shift for concentration of chlorophyll from October 2011 to March 2015. (D) shows the results of the time shift for sea surface temperature from October 2011 to March 2015.

The space-shift experiment showed that observed turtle locations were associated with intermediate SST and Chl-a values relative to the minimum and maximum values available in adjacent spatial alternatives (Fig. 6A–D). Observed conditions differed significantly from both lower and higher extremes, suggesting that oceanic foragers used environmental conditions within the central part of the local thermal–trophic gradient rather than consistently occupying the coldest, warmest, least productive or most productive available areas (LME F = 345.6 for Chl-a; F = 44.5 for SST; both p < 0.001, Table S6; Fig. 6 C&D).

**Figure 6.**
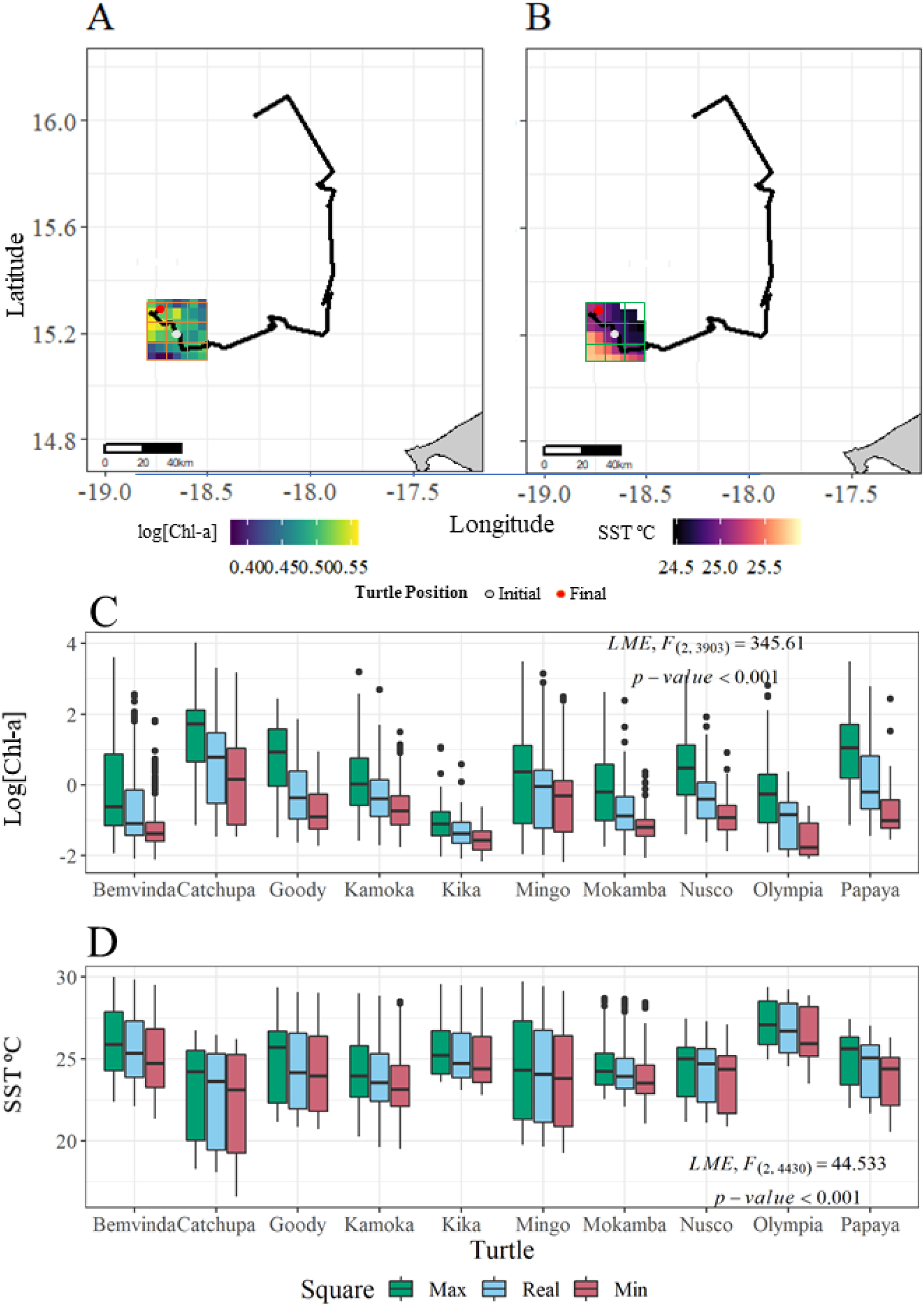
Spatial shift experiments. A & B the spatial shift experiment Nusco track for two weeks, from 28^th^ November to 12^th^ December 2011, showing the [Chl-a] and SST polygon for the last day of the track, the gray dot is the initial position of the Mokamba on 11^th^ December and the red one is the first location of the Makamba on 12^th^ December. C-D) Boxplot with the experimental results for every day that turtles transmitted: the concentration of chlorophyll-a is shown in (C) while sea surface temperature in (D). In green is the polygon with the maximum values, in red is the polygon with the minimum values and in blue is the polygon for the conditions experienced by the turtles.

#### Remigration

Because all female turtles were PIT tagged, we were able to infer fidelity and remigration intervals based on detection on the nesting beaches in later years. All three female turtles tracked from Boa Vista in 2011 were detected nesting again on the same island in 2013 (Bolacha and Nusco) and 2018 (Goody), indicating minimum remigration periods of 2 years. One female turtle tracked from Sal (Papaya) in 2011 was observed nesting on the same island again in 2017. None of the female turtles were detected nesting within 1 year of being tagged on their nesting island. On the other hand, the migratory male was tracked for 738 days from July 2013. He left Cabo Verde on 20 July 2013, towards the foraging area and returned at the end of November 2013, left again in December 2013 towards the foraging area and came back to Cabo Verde in August 2014, and June 2015. When transmission ended his trajectory suggests he was swimming back to Cabo Verde (Fig. 4D). This turtle likely contributed to the mating season three years in a row.

## Discussion

Identifying the specific environmental needs of marine megafauna during different life stages is key to understanding the habitat-specific movements that shape their behaviour. Such understanding also helps define species’ environmental niches and predict future distributions. Here, we focused on loggerhead sea turtles nesting in the Cabo Verde archipelago. In the breeding habitat, we identified two contrasting movement patterns, where female turtles nesting on western islands, that lack shallow coastal areas, showed wide-ranging movements, whereas males and females breeding on eastern islands with extensive shallow zones tended to remain nearshore. Two of the three tagged males remained resident within the breeding habitat throughout the tracking period. In the migratory corridor, all migratory turtles (nine females and one male) exhibited directional movements toward the west coast of Africa, independently of the environmental variables tested. In the foraging habitat, neritic and oceanic foragers experienced distinct SST and chlorophyll-a (Chl-a) regimes. Using two virtual experiments, we showed that turtle movements were constrained by a negative correlation between SST and Chl-a concentration. We suggest that daily displacements at oceanic foraging grounds reflect an optimal foraging strategy that allows turtles to track prey, reduce predation risk, and maintain an active metabolism.

At breeding sites, females typically rest in shallow waters, where they can optimise dive durations and access warm patches that accelerate egg development (Mortimer and Portier 1989, Hays et al. 2000, Schofield et al. 2009). In Cabo Verde, females nesting on islands with such habitats (<20 m isobath, generally on the eastern side of the archipelago) remained within 2 km of the coast. In contrast, on islands where nearshore habitats drop abruptly (>1000 m isobath within 10 km from shore), females undertook continuous large loops up to 360 km during each inter-nesting period. Similar wide-ranging behaviour occurs in leatherbacks nesting on islands with narrow peri-insular shelves that descend rapidly to 1000 m (Georges et al. 2007, Byrne et al. 2009). Leatherbacks are believed to forage during this period (Myers and Hays 2006). In loggerheads, long-distance “foray” movements may be associated with feeding (BBC, Animal with Cameras), but also with the capacity of females to explore alternative nesting habitats (Blumenthal et al. 2006, Rees et al. 2010, Schofield et al. 2010c, Hart et al. 2013). Regardless of the underlying cause, this behavioural dichotomy likely reflects individual differences in the balance between energetic costs and reproductive benefits. Wide-ranging turtles may reduce predator encounters or exploit transient prey, while more sedentary turtles minimise energetic costs by remaining in optimal shallow areas (Bonnet et al. 1998), where water temperature supports efficient egg incubation (Schofield et al. 2009).

After breeding in Cabo Verde, one male and ten females migrated to non-breeding areas, showing strongly directional movement toward foraging habitats, as also reported for other loggerhead populations in the Mediterranean and North Pacific (Schofield et al. 2010b, Abecassis et al. 2013, Stokes et al. 2015, Almpanidou et al. 2019). The male’s migration differed from that of females, departing earlier and following a warmer thermal corridor, consistent with observations from the Mediterranean, where males experience higher sea surface temperatures during migration (Schofield et al. 2010b, Almpanidou et al. 2019). We found no sex differences in daily displacement or directional persistence, unlike populations in Japan and the Caribbean Sea where males and females follow distinct migratory routes (Sakamoto et al. 1997, Van Dam et al. 2008). In their migratory corridors, turtles crossed low-productivity regions, supporting the idea that they feed only opportunistically during transit (Luschi et al. 1998, Jonsen et al. 2005, Godley et al. 2008, Dujon et al. 2017). Interestingly, migration from Cabo Verde occurred against prevailing ocean currents, requiring constant directional correction as suggested by turning angle analyses. This behaviour likely imposes high energetic costs and indicates a strong navigational control rather than passive drift, consistent with patterns described for Mediterranean loggerheads (Hays et al. 2014).

In the foraging habitat, we observed both oceanic and coastal (neritic) strategies in males and females, as documented in Japan, the southeastern United States, and Greece (Hatase et al. 2002, Reich et al. 2009, Schofield et al. 2010c). Notably, neritic males remained resident in Cabo Verde waters, whereas the neritic female migrated to Sierra Leone, approximately 1,280 km from the nesting area. Earlier telemetry studies estimated that about 30% of turtles from Boa Vista were neritic foragers (Hawkes et al. 2006, Eder et al. 2012, Varo-Cruz et al. 2013). However, a larger stable isotope study found that neritic foragers represent only about 20% of individuals, mainly from eastern islands and particularly Boa Vista (Cameron et al. 2019). Our findings, that include turtles from more islands, support the view that oceanic foraging predominates across the population. Oceanic individuals followed large looping trajectories typical of deep-water foragers, similar to loggerheads from Japan, the Arabian Sea, and the Mediterranean (Hatase et al. 2002, Rees et al. 2010, Schofield et al. 2010b).

Neritic and oceanic females did not return for at least two years, whereas the migratory male returned multiple times to Cabo Verde. Males are known to visit breeding areas up to 2.6 times more often than females, resulting in an operational sex ratio closer to parity than the female-biased hatchling ratio (Limpus 1993, James et al. 2005, Schofield et al. 2017). Our data suggest two male strategies that may maintain this balance: resident males reduce migration costs while ensuring frequent mating opportunities, whereas oceanic males range widely, potentially increasing encounters with females from multiple islands. This far-reaching movement pattern supports the male-mediated gene flow previously reported for the Cabo Verde population (Stiebens et al. 2013, Baltazar-Soares et al. 2020).

Animals invest in efficient foraging strategies, a key requirement for capital breeders such as sea turtles that must replenish energy stores between breeding seasons (Stephens et al. 2009, Plot 2013, Perrault and Stacy 2018). Turtles foraging in neritic areas are likely to target energy-rich benthic prey but face higher predation risk (Heithaus 2013). In contrast, oceanic foragers must travel long distances across patchy prey fields, sometimes hundreds of kilometres, to locate sufficient food (Kobayashi et al. 2008, Schofield et al. 2010b). Our results indicate that oceanic foragers exploit an optimal foraging niche defined by the negative correlation between SST and Chl-a concentration (Kobayashi et al. 2008, Howell et al. 2015). Noteworthy, Chl-a should be interpreted as an indirect proxy for potential prey availability rather than a direct measure of turtle food resources. The strength and timing of the relationship between phytoplankton biomass and gelatinous or other prey may vary across regions, seasons and trophic pathways. Therefore, our results identify environmental conditions associated with turtle space use, but do not directly demonstrate prey capture or diet composition. Because Chl-a reflects phytoplankton biomass, it represents the foundation of the food web and can indicate areas where gelatinous prey such as jellyfish may become more abundant (Thompson et al. 2012, Colella et al. 2016, Polovina et al. 2017). Along the northwest African coast, upwelling events create phytoplankton blooms that link cold, nutrient-rich waters with high productivity, reinforcing the observed temperature–Chl-a trade-off (Menna 2016). Our time-shift experiment shows that this relationship not only shapes spatial distribution but also constrains the timing of arrival at foraging grounds, indicating a fine-scale match between movement and environmental cycles. Similarly, seasonal habitats of North Pacific loggerheads were defined by overlapping SST and Chl-a conditions, though at slightly lower SST ranges (Kobayashi et al. 2008).

The spatial-shift experiment further supports this interpretation: turtles were associated with intermediate Chl-a and SST values relative to local environmental alternatives. This pattern is consistent with use of a restricted environmental window, although it should not be interpreted as direct proof of active selection without additional null-track comparisons (Polovina et al. 2001, Kobayashi et al. 2008, Block et al. 2011, Howell et al. 2015). Intermediate productivity likely reflects the most profitable foraging conditions, balancing prey availability and search costs (Olson et al. 1994, Polovina et al. 2001). Similar strategies occur in other marine predators, including albacore tuna that forage along North Pacific fronts and fin whales whose distributions track productivity gradients in the Mediterranean (Polovina et al. 2001, Cotte et al. 2010). Together, these findings reveal that the timing and location of foraging depend on environmental thresholds that guide turtles toward prey-rich yet thermally suitable waters while avoiding energetically costly extremes. Our study therefore identifies a mechanistic proxy that can inform models predicting turtle movements under changing ocean conditions.

Overall, by examining movements across multiple habitat types, we show that adult loggerhead turtles breeding in Cabo Verde respond to distinct environmental drivers that shape their spatial behaviour throughout the annual cycle. Although movements are complex, turtles appear to use environmental conditions that may reduce energetic costs while maintaining access to productive foraging areas. This behavioural plasticity demonstrates how physical and biological ocean features interact with evolved adaptive processes to influence movement decisions. As oceans warm and productivity regimes shift, the balance between thermal and trophic constraints may redefine foraging suitability, migration timing, and reproductive connectivity. Our findings thus contribute to understanding how marine megafauna integrate environmental cues into movement decisions and provide an ecological foundation for predicting responses of this endangered population to climate-driven ocean change

## Acknowledgments

The authors would like to thank members of all NGOs involved in this study for their support in the field, Fundação Tartaruga, Associação Projecto Vitó, Associação Projeto Biodiversidade, as well as the team from the Instituto do Mar.

## Funding

This work was funded by grants from the National Geographic (NGS-13; NGS-59158R-19 to C.E.) grants. Field seasons were further supported by UK Research and Innovation Natural Environment Research Council (NERC, NE/V001469/1, NE/X012077/1). P.R.T is funded by the Mexican National Council for Science and Technology (CONACYT).

## Authors’ contributions

Project Conception: C.E, V.A.S, P.R.T; Field work: V.A.S, C.R., H.D., A.T., S.C., C.E; Data Analyses and guidance: P.R.T, G.S., G.C.H, C.E; Drafting of the manuscript: P.R.T.; Review: All authors.

## Supplementary material

**Table S1.** Attachment details of tracked turtles.

| Turtle Name | PTT | S | CCL/ CCW x [cm] | Manufacturer | Deployment date | Deployment location | Days transmitted | Total displacement (km) | Min/Max Displacement per day (km) | Departure date from breeding area | Foraging Type |
| --- | --- | --- | --- | --- | --- | --- | --- | --- | --- | --- | --- |
| **Nusco** | 102257 | F | 85/76 | SMRU | 26.07.2011 | Lacacao, Boa Vista | 311 | 4,428 | 3.6/29.12 | 01/09/2011 | Oceanic |
| **Bolacha** | 102258 | F | 90/80.2 | SMRU | 26.07.2011 | Lacacao, Boa Vista | 399 | 1,770 | 2.68/13.16 | 29/09/2011 | Neritic/<br>Migrant |
| **Goody** | 108522 | F | 85.3/79.4 | SMRU | 23.07.2011 | Lacacao, Boa Vista | 260 | 4,868 | 9.42/28.84 | 10/09/2011 | Oceanic |
| **Catchupa** | 108523 | F | 77/73 | SMRU | 22.09.2011 | Serra Negra, Sal | 149 | 2,743 | 3.02/25.35 | 29/09/2011 | Oceanic |
| **Manga** | 108524 | F | 82/78.5 | SMRU | 24.09.2011 | Serra Negra, Sal | 60 | 28* | 1.26/1.83 | NA | NA |
| **Papaya** | 108525 | F | 79/75 | SMRU | 23.09.2011 | Serra Negra, Sal | 124 | 2,887 | 2.6/31.53 | 24/09/2011 | Oceanic |
| **Nhanha** | 118292 | F | 79/70 | Wildlife | 07.08.2012 | S. Catarina, Fogo | 27 | 791 | NA/NA** | NA | Oceanic |
| **Bemvinda** | 118293 | F | 87.5/83 | Wildlife | 31.07.2012 | Lazareto, S. Vicente | 1109 | 17,073 | 1.66/38.61 | 03/09/2012 | Oceanic |
| **Kika** | 11829<br>4 | F | 77/77 | Wildlife | 01.08.2012 | Lazareto,<br>S. Vicente | 867 | 9,912 | 2.95/31.97 | 31/08/2012 | Oceanic |
| **Kamoka** | 11874<br>2 | F | 88/83 | Sirtrack | 06.08.2012 | S.<br>Catarina,<br>Fogo | 944 | 16,035 | 11.45/25.4 | 04/09/2012 | Oceanic |
| **Olympia** | 11874<br>3 | F | 77/75 | Sirtrack | 01.08.2012 | Lazareto,<br>S. Vicente | 156 | 2,802 | 10.34/32.72 | 01/10/2012 | Oceanic |
| **Mokamba** | 11874<br>4 | F | 80/79 | Sirtrack | 08.08.2012 | S.<br>Catarina,<br>Fogo | 400 | 8,033 | 13.91/29.61 | 24/09/2012 | Oceanic |
| **Fra** | 13007<br>5 | M | 105/94 | Wildlife | 30.06.2013 | Salrei,<br>Boa Vista | 244 | 1,349 | 5.71/6.36 | Didn't<br>migrate | Neritic/<br>Resident |
| **Mingo** | 13007<br>6 | M | 81/79 | Wildlife | 30.06.2013 | Salrei,<br>Boa Vista | 738 | 12, 651 | 9.66/24.24 | 20/07/2013 | Oceanic |
| **Ze** | 13007<br>7 | M | 86/81 | Wildlife | 20.07.2013 | Salrei,<br>Boa Vista | 60 | 377 | 3.16/4.61 | Didn't<br>migrate | Neritic/<br>Resident |

\*Total displacement from the first 30 days when the turtle was in the breeding ground, then only rare locations were transmitted and no further displacement was estimated to avoid possible bias. \*\*An internal tag mis-function affected the timing but reported the location correctly, as such daily displacement could not be estimated.

**Table S3.** Statistical test between phases.

|  | *Breeding* | *Travelling* | *Foraging* | *Test* | *Statistics* | *Tukey test* |
| --- | --- | --- | --- | --- | --- | --- |
| Chl-a (mg m<sup>-3</sup>) | -1.692<br>SE ± 0.39 | -1.417 SE ± 1.13 | -0.394<br>SE ± 1.02 | LME | F<sub>(2,111)</sub> = 13.76,<br>p < 0.001 | Z<sub>(Breeding- migration )</sub> = -5.47, p < 0.001,<br>Z<sub>(Breeding- foraging)</sub> = -6.353, p < 0.001,<br>Z<sub>(migration- foraging)</sub> = 297.2, p = 0.471 |
| SST (°C) | 25.829<br>SE ± 0.87 | 28.107 SE ± 0.710 | 24.818<br>SE ± 2.621 | LME | F<sub>(2,111)</sub> = 7.207,<br>p < 0.001 | Z<sub>(Breeding- migration )</sub> = 2.128, p = 0.081,<br>Z<sub>(Breeding- foraging)</sub> = 1.353, p = 0.357,<br>Z<sub>(migration- foraging)</sub> = 3.696, p < 0.001 |
| Bathymetry (m) | 818.71<br>SE ± 995.18 | 3062.99 SE ± 394 | 2916.21<br>SE ± 1231.34 | LME | F<sub>(2,111)</sub> = 22.57,<br>p < 0.001 | Z<sub>(Breeding- migration )</sub> = -5.47, p < 0.001, Z<sub>(Breeding- foraging)</sub> = -6.353, p < 0.001, Z<sub>(migration- foraging)</sub> = 297.2, p = 0.47 |
| Displacement (km/day) | 10.31 SE<br>± 8.40 | 28.31 km SE<br>± 3.61 | 17.25 km<br>SE ± 6.32 | LME | F<sub>(2,111)</sub> = 7.678,<br>p = 0.004 | Z<sub>(Breeding- migration )</sub> = 17.45, p < 0.001,<br>Z<sub>(Breeding- foraging)</sub> = -5.710, p = 0.005, Z<sub>(migration- foraging)</sub> = 11.74, p < 0.001 |
| Turning angle | Rayleigh<br>test, F =<br>0.106,<br>p < 0.001 | Rayleigh<br>test, F =<br>0.359,<br>p < 0.001 | Rayleigh<br>test, F =<br>0.333,<br>p < 0.001 | Circular | F<sub>(2,14181)</sub> = 3.85,<br>p = 0.021 |  |

**Table S4.**
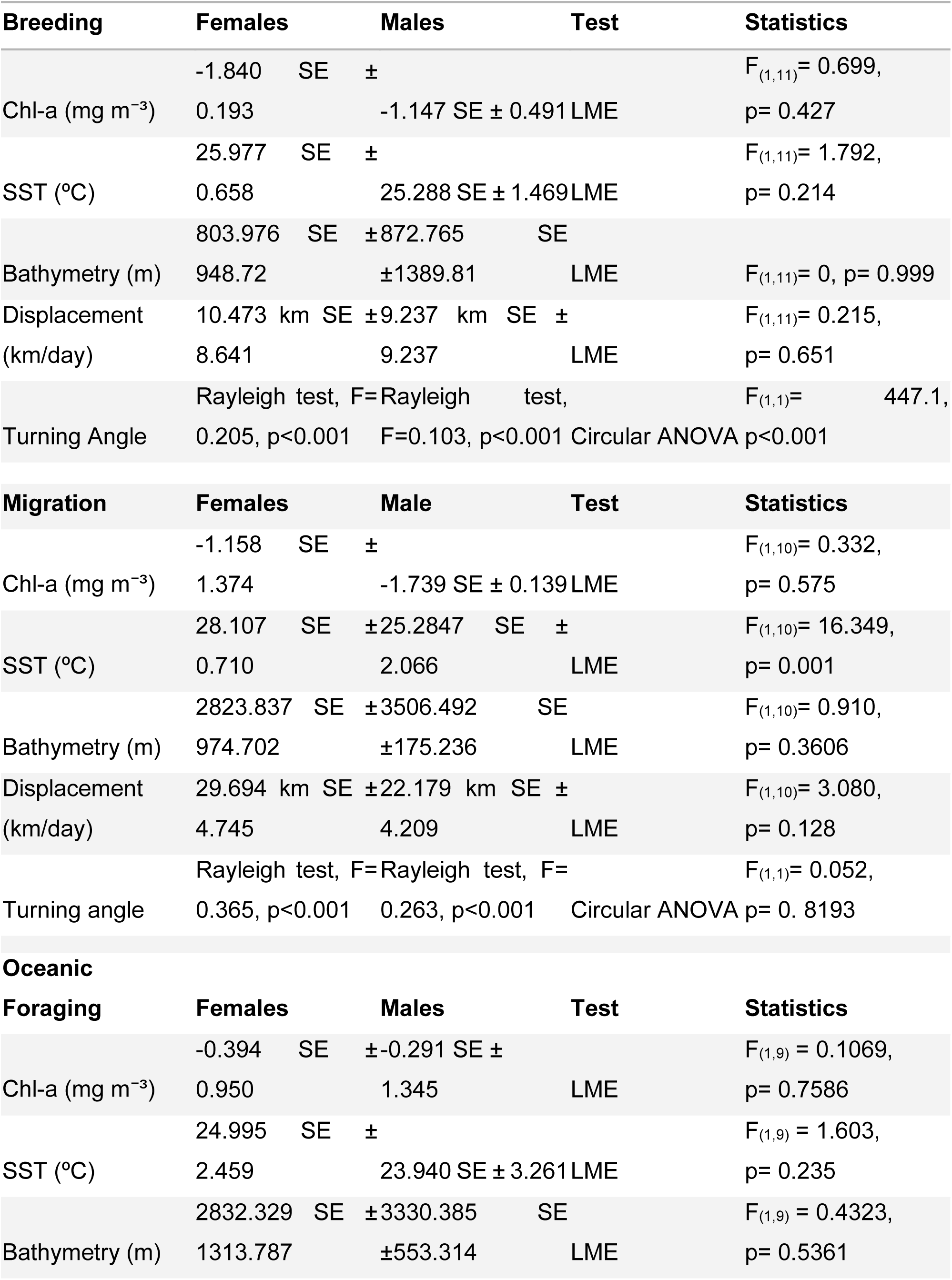
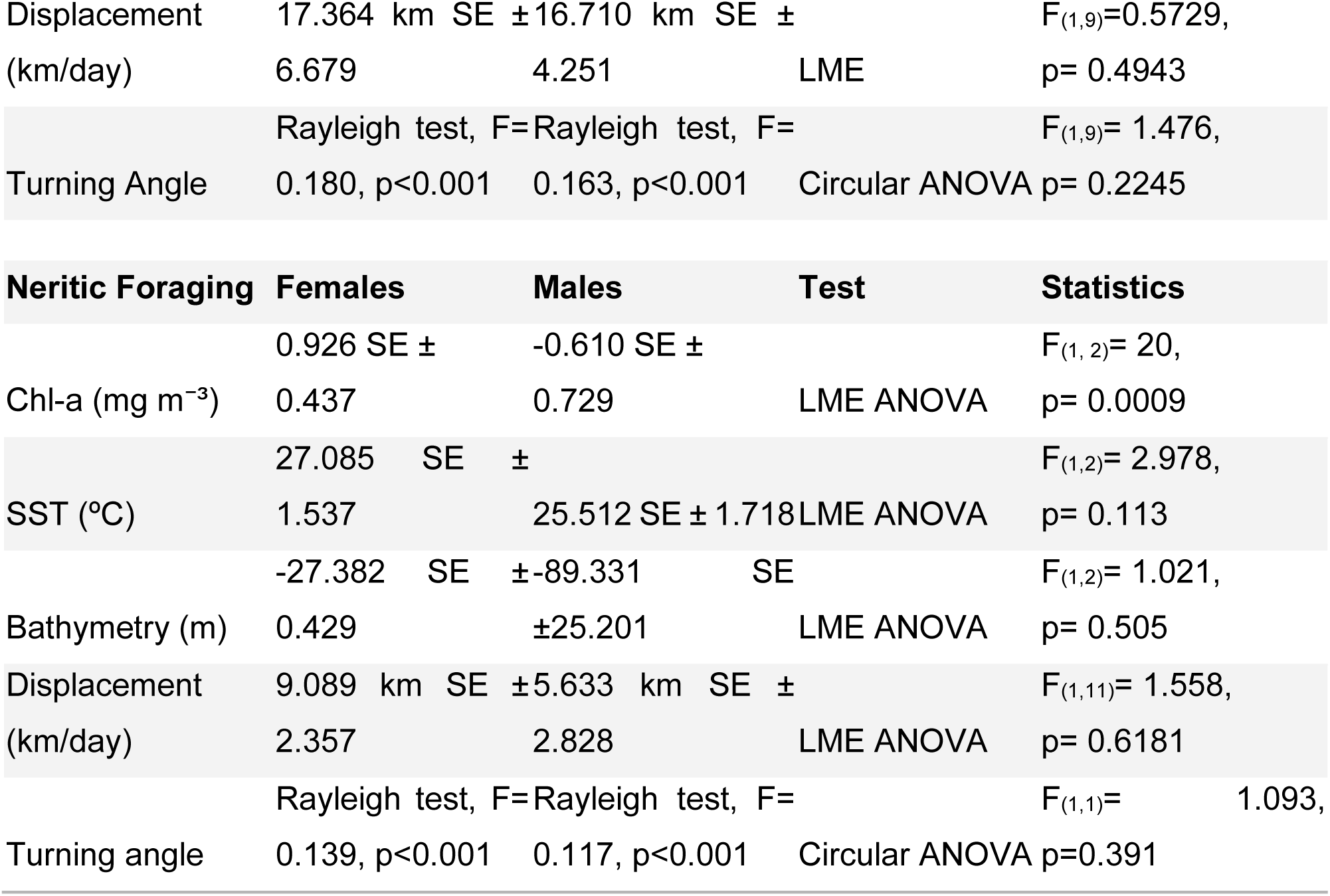
Statistical test between sexes in each phase.

**Table S5.**
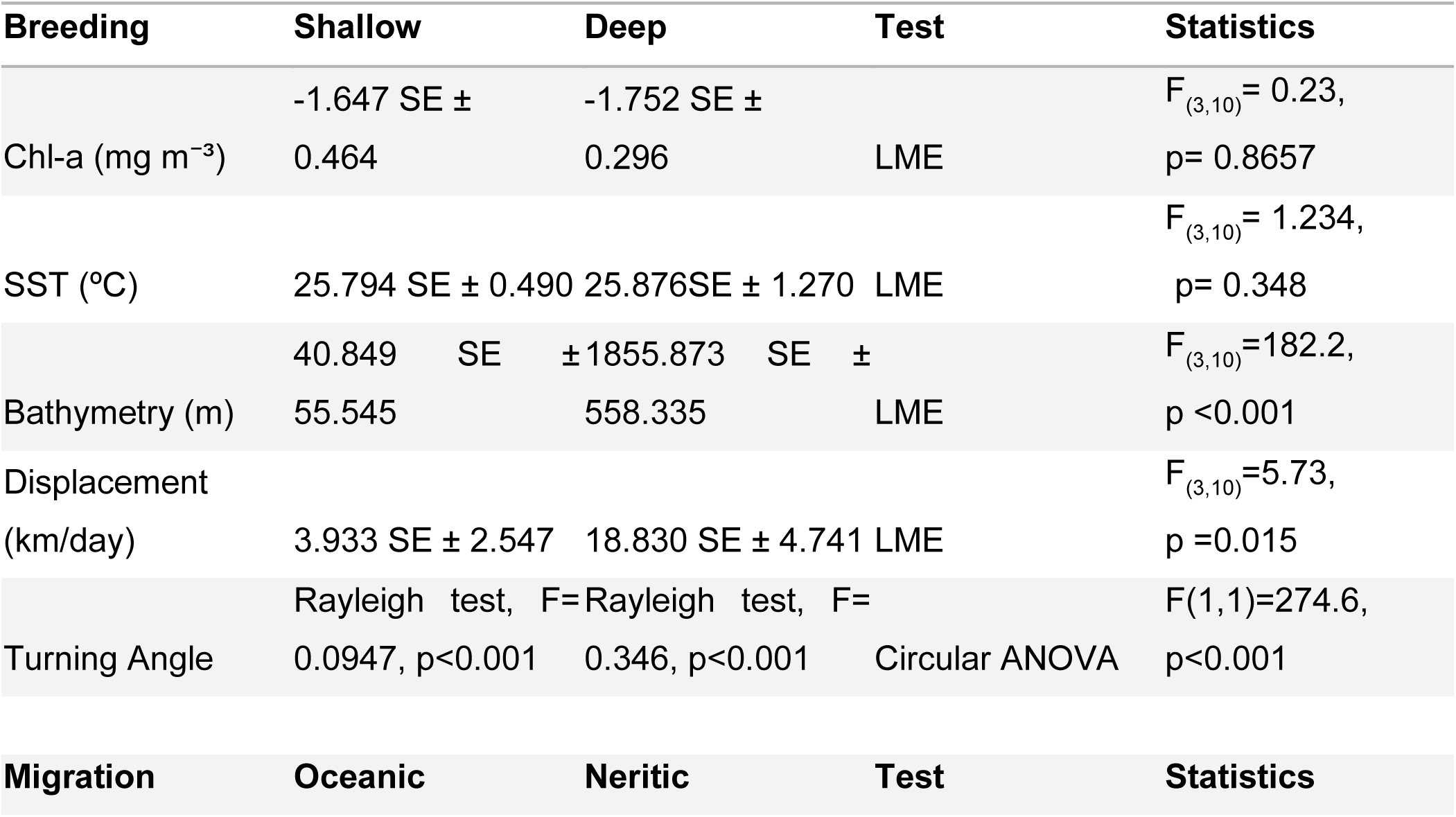
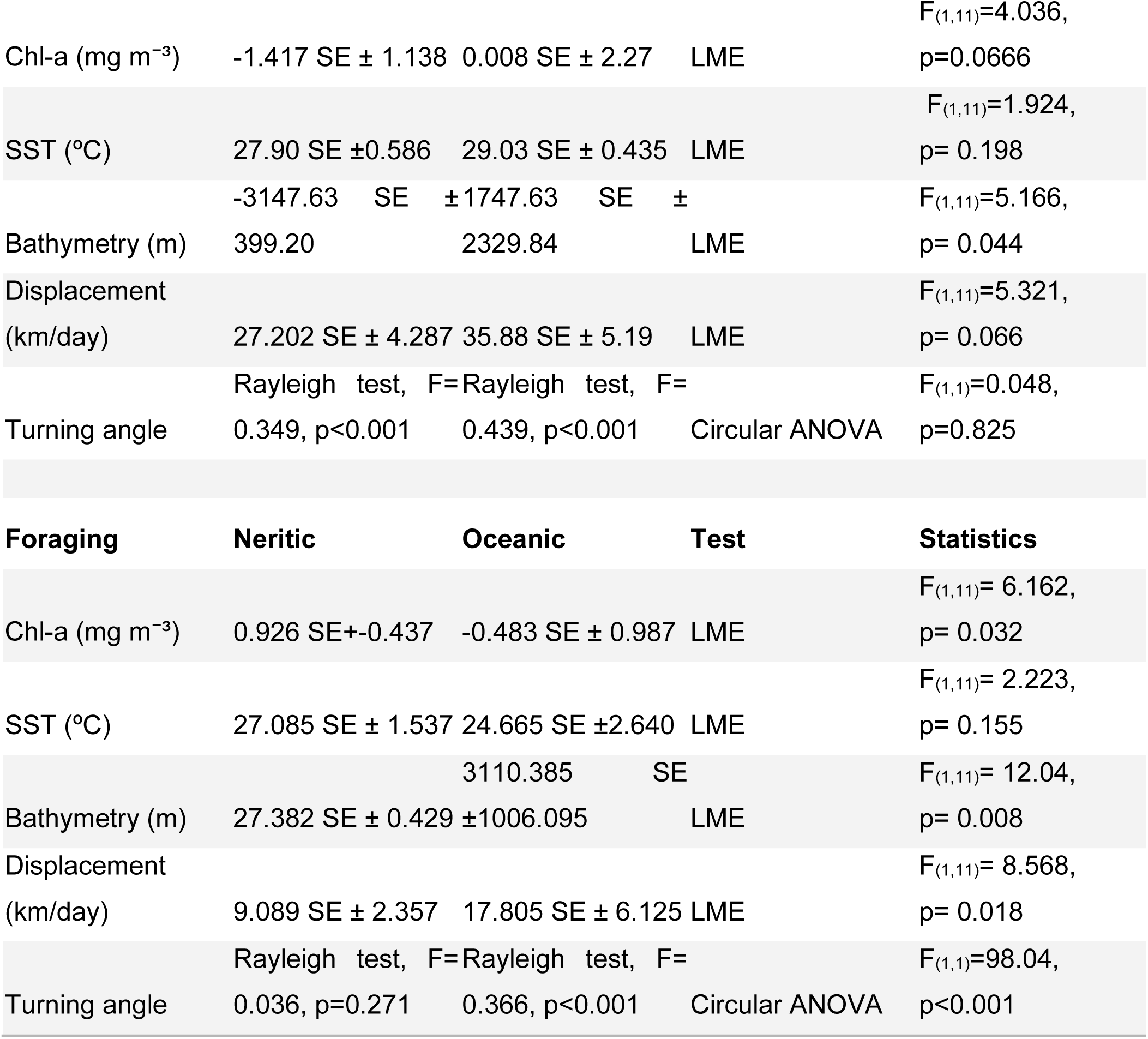
Statistical tests between behaviours in each phase.

**Table S6.** Statistical test for the time and spatial shift experiment.

| Time shift experiment | Previous | Real | After | Test | statistics | Tukey test |
| --- | --- | --- | --- | --- | --- | --- |
| Chl-a (mg m<sup>-3</sup>) | -0.765,<br>SE±<br>0.763 | -0.424<br>SE ±<br>1.053 | 0.246<br>SE±<br>0.976 | LME | F<sub>(3, 240)</sub>=9.18, p<<br>0.001 | Z (Previous-After)=-4.190, p < 0.001,<br>Z(Real-After)= -1.315, p = 0.385,<br>Z (Real-Previous)= 2.875, p = 0.0112 |
| SST (°C) | 25.309<br>SE±<br>2.667 | 24.517<br>SE±<br>2.557 | 23.402<br>SE±<br>2.522 | LME | F<sub>(3,276)</sub> =27.45, p<<br>0.001, | Z(Previous-After)=-5.494, p < 0.001,<br>Z(Real-After)= 2.581, p = 0.0265,<br>Z(Real-Previous)= -2.913, p = 0.010 |
| Spatial shift experiment | Max | Real | Min | Test | Statistics | Tukey test |
| Chl-a (mg m<sup>-3</sup>) | 0.130<br>SE ±<br>1.256 | -0.424<br>SE ±<br>1.053 | -0.860<br>SE ±<br>0.856 | LME | F<sub>(2,3903)</sub> = 345.61,<br>p<0.001 | Z (Min-Max)=-26.23, p < 0.001,<br>Z(Real-Max)= -14.70, p < 0.001,<br>Z (Real-Min)= 11.53, p < 0.001 |
| SST (°C) | 24.867<br>SE<br>±2.571 | 24.517<br>SE±<br>2.557 | 24.091<br>SE ±<br>2.539 | LME | F<sub>(2,4430)</sub> = 44.533,<br>p<0.001 | Z(Min-Max)=-9.423, p < 0.001,<br>Z (Real-Max)= -4.257, p < 0.001,<br>Z (Real-Min)= 5.166, p < 0.001 |

**Figure S1.**
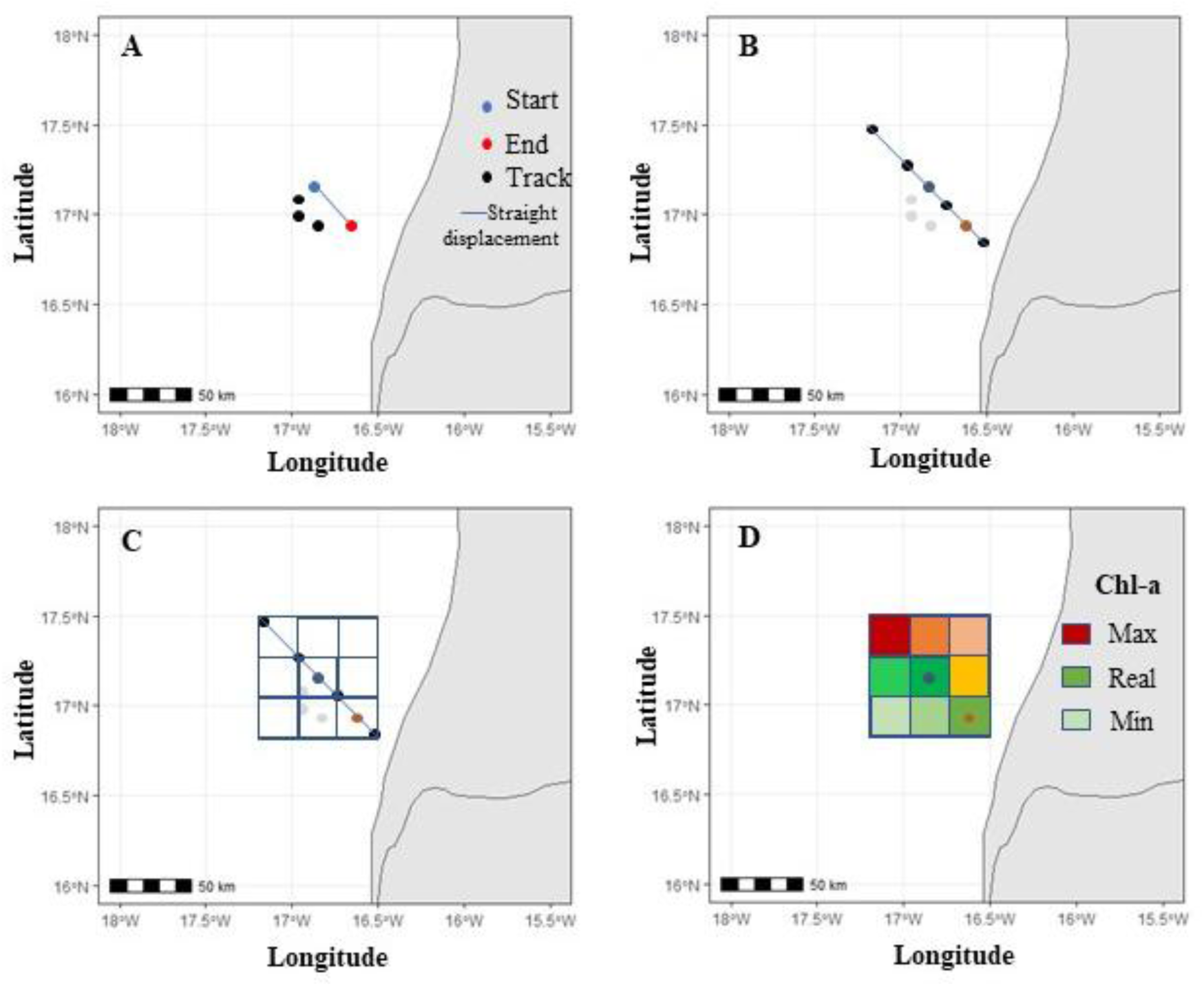
Spatial shift experiment. A) A turtle track over two days, the start is the initial position of the turtle and the end is the first location of the turtle in the following day. The straight line shows the shortest displacement between the start and the end. B) From the straight line displacement, the side length of the polygons are calculated using Pythagoras theorem (*h^2^= 2a^2^*, where *h* is the straight line displacement, and a the side of the polygon) and the vertex positions of each polygon are estimated, the vertex of the central polygon are located in the middle of the straight line displacement (black dots) from the initial position of the turtle and the vertexes of the polygons in the diagonal are located at 1.5 times of the straight line displacement (yellow dots). C) Projection of the 9 polygons. D) The mean Chlorophyll-a concentrations are extracted for each polygon and then ranked from the highest (maximizing) to the lowest (minimizing).

**Figure S2.**
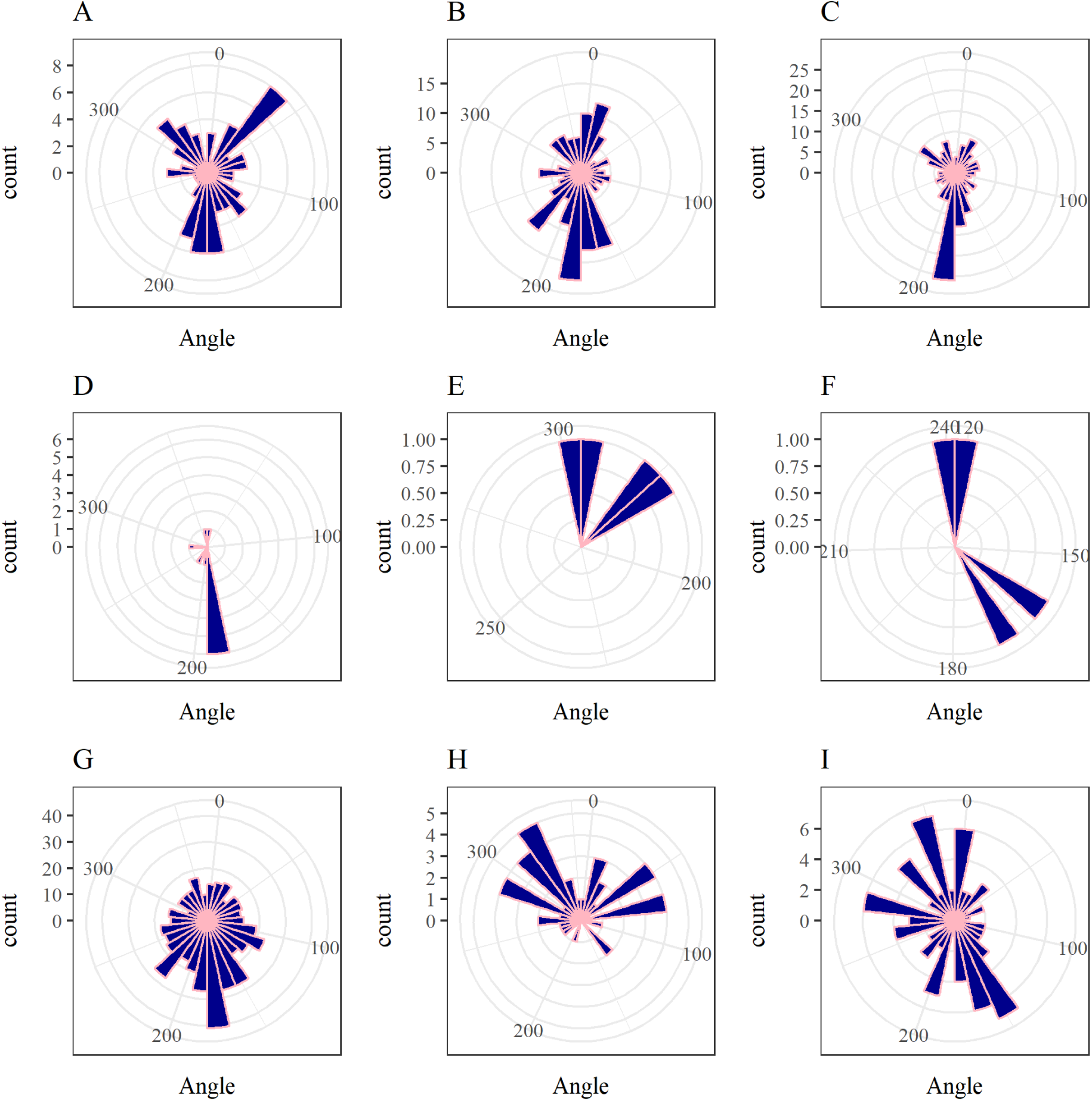
Turning angles histograms from the movements during the breeding habitat. From Boa Vista: A) Bolacha, B) Nusco, C) Goody. From Sal: D) Catchupa, E) Manga, F) Papaya. Males from Boa Vista: G) Fra, H) Mingo, I) Ze.

**Figure S3.**
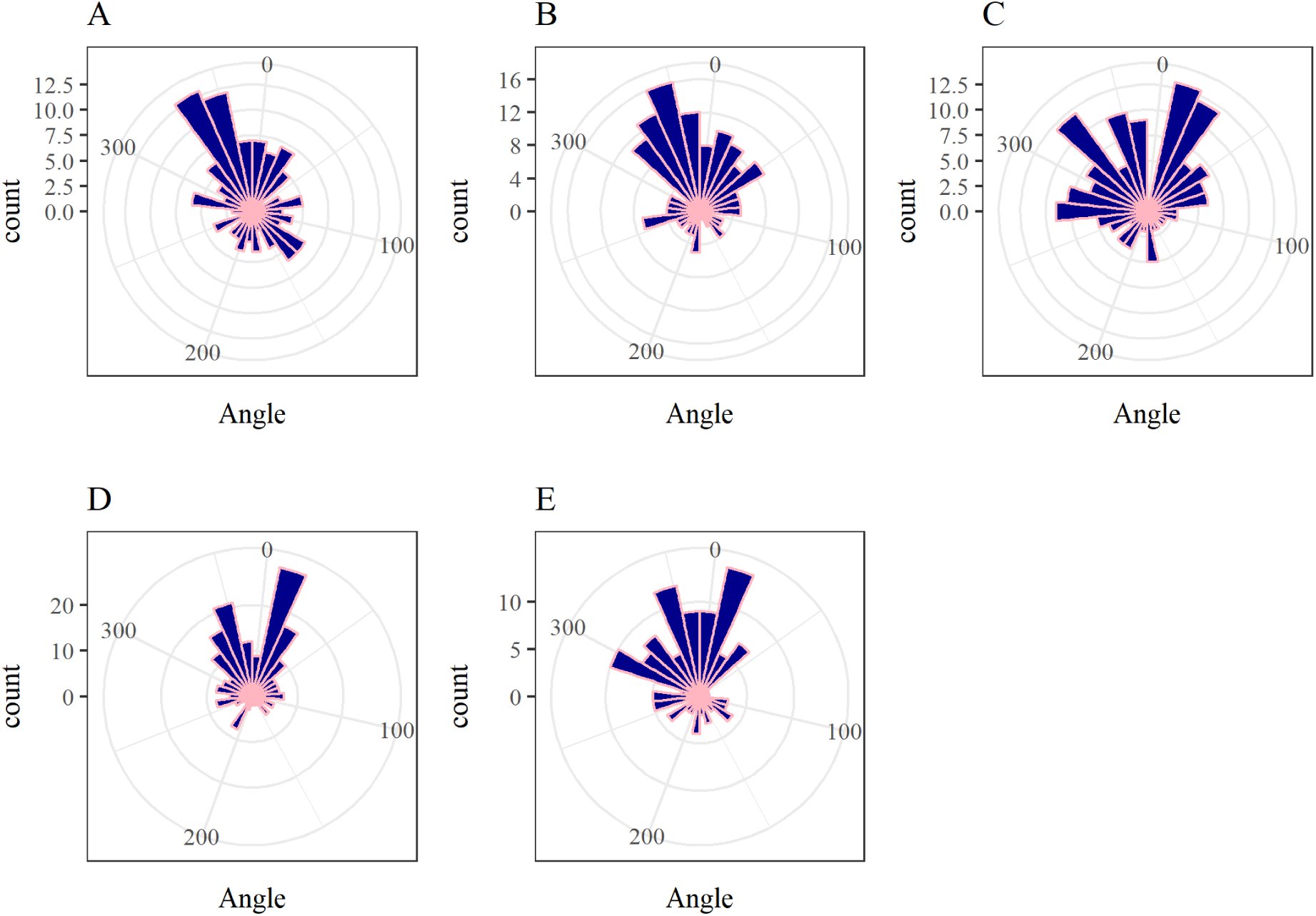
Turning angles histograms from the movements during breeding habitat. From São Vicente: A) Bemvinda, B) Kika, C) Olympia. From Fogo: D) Mokamba, E) Kamoka.

**Figure S4.**
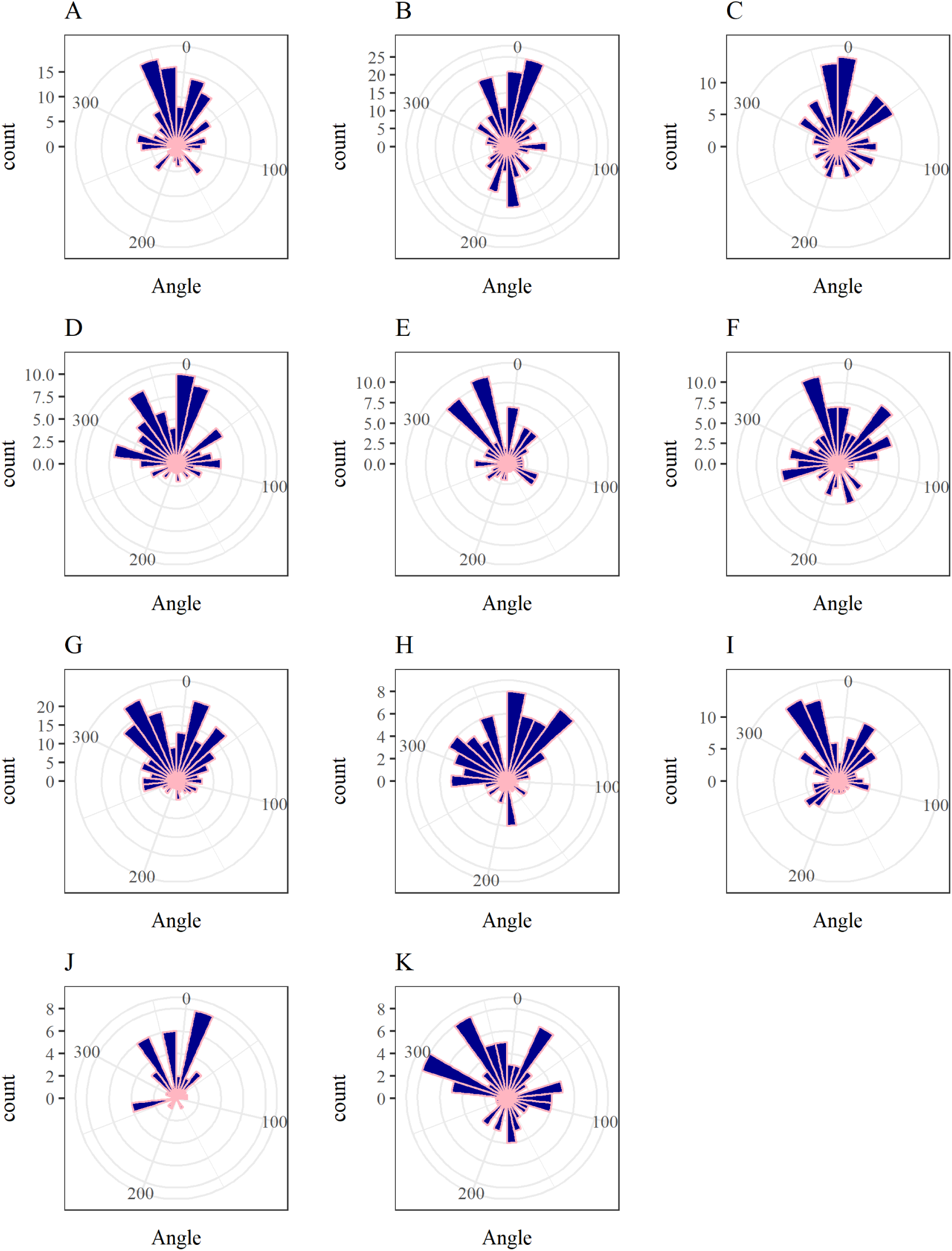
Turning angles histograms from the movements during the migrating route. From Boa Vista: A) Bolacha (Neritic), B) Nusco, C) Goody. From Sal: D) Catchupa and E) Papaya. From São Vicente: F) Bemvinda, G) Kika, H) Olympia. From Fogo: I) Mokamba, J) Kamoka. The male from Boa Vista: K) Mingo.

**Figure S5.**
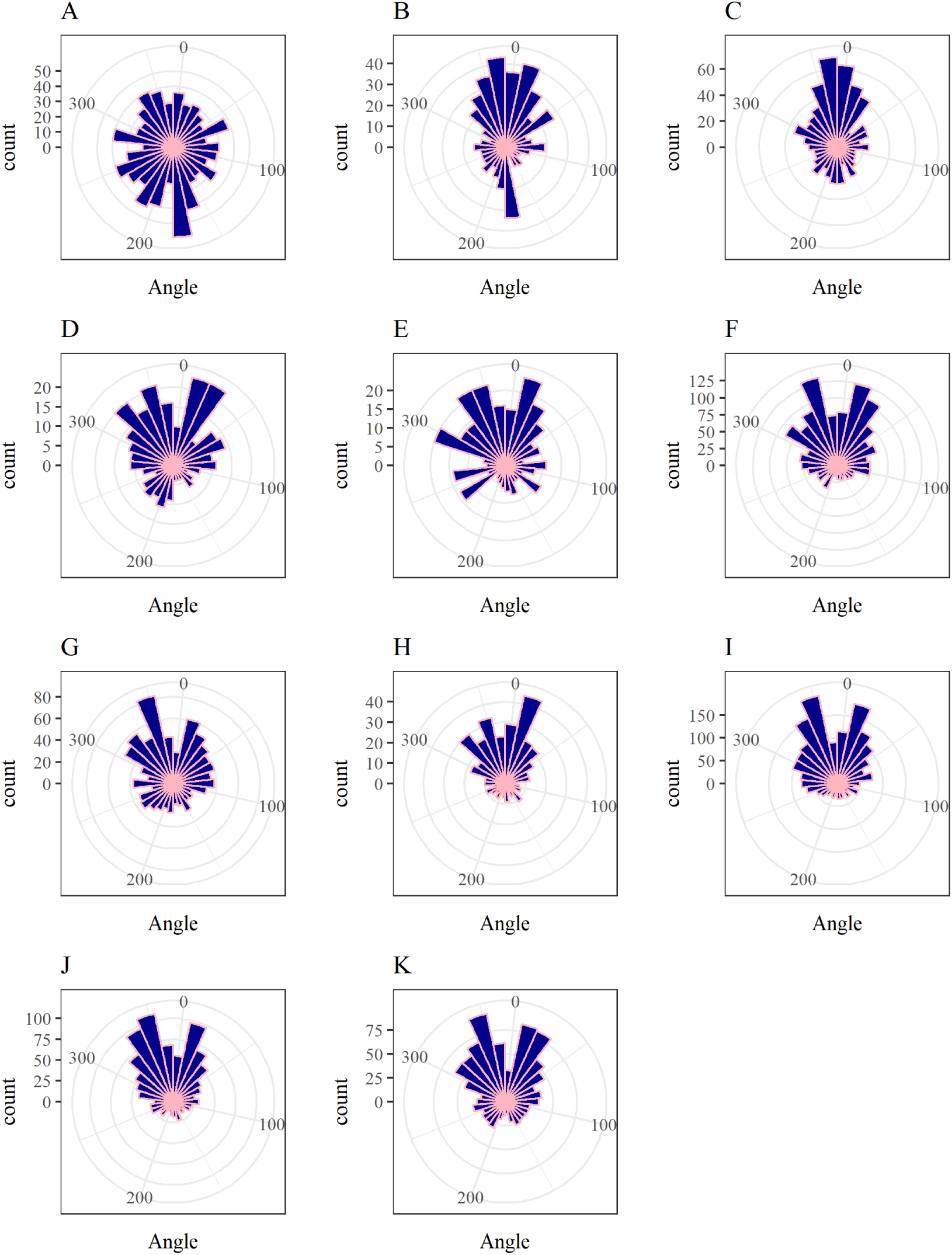
Turning angles histograms from the movements during the migrating route. From Boa Vista: A) Bolacha (Neritic), B) Nusco, C) Goody. From Sal: D) Catchupa and E) Papaya. From São Vicente: F) Bemvinda, G) Kika, H) Olympia. From Fogo: I) Mokamba, J) Kamoka. The male from Boa Vista: K) Mingo.

**Figure S6.**
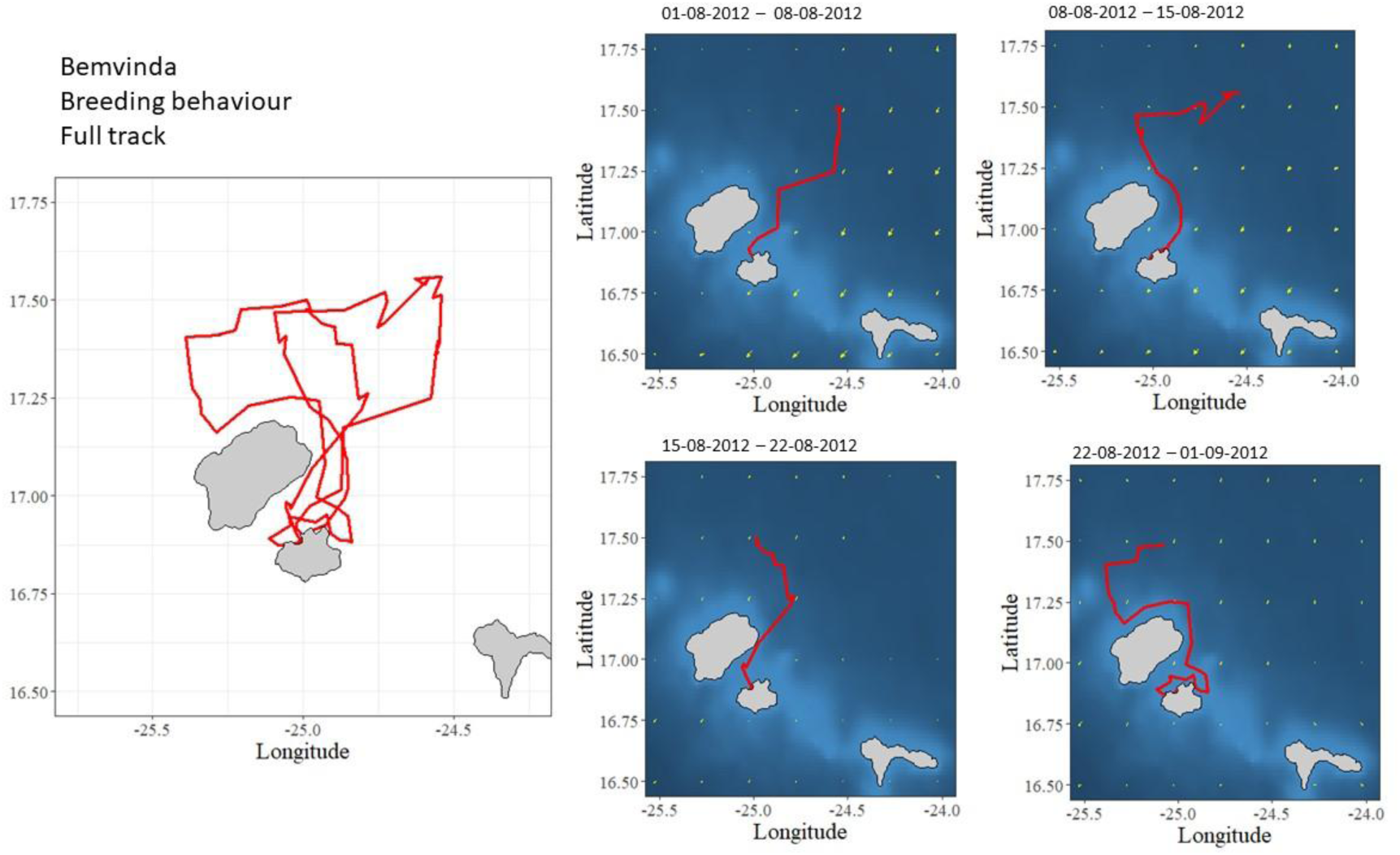
Bemvinda breeding behaviour. A) Full track, two loops anticlockwise. B)Left the island going north against the currents between 1^st^ and 8^th^ August 2012. C)Come back to the nesting beach following the currents 8^th^ and 15^th^ August 2012. D) Left the island going north against the currents 15^th^ and 22^nd^ August 2012, E) Come back to the nesting beach following the currents 22^nd^ August and 1^st^ September 2012.

## Notes

### Competing Interest Statement

The authors have declared no competing interest.

### Summary of Updates

This version is a correction linked to a typo in the affiliation of the author Herculano A Dinis, Associacao Projeto Vito, Sao Filipe, Fogo, Cabo Verde.

